# Modulation of tissue growth heterogeneity by responses to mechanical stress

**DOI:** 10.1101/425355

**Authors:** Antoine Fruleux, Arezki Boudaoud

## Abstract

Morphogenesis often yields organs with robust size and shapes, whereas cell growth and deformation feature significant spatio-temporal variability. Here, we investigate whether tissue responses to mechanical signals contribute to resolve this apparent paradox. We built a model of growing tissues made of fiber-like material, which may account for the cytoskeleton, polar cell-cell adhesion, or the extracellular matrix in animals, and for the cell wall in plants. We considered the synthesis and remodeling of this material, as well as the modulation of synthesis by isotropic and anisotropic response to mechanical stress. Formally, our model describes an expanding, mechanoresponsive, nematic, and active fluid. We show that mechanical responses buffer localized perturbations, with two possible regimes - hypo-responsive and hyper-responsive, and the transition between the two corresponds to a minimum value of the relaxation time. Whereas robustness of shapes suggests that growth fluctuations are confined to small scales, our model yields growth fluctuations that have long-range correlations. This indicates that growth fluctuations are a significant source of heterogeneity in development. Nevertheless, we find that mechanical responses may dampen such fluctuations, with a specific magnitude of anisotropic response that minimizes heterogeneity of tissue contours. We finally discuss how our predictions might apply to the development of plants and animals. Altogether, our results call for the systematic quantification of fluctuations in growing tissues.

---

Variability has emerged as an inherent feature of many biological systems (1, 2), spanning molecular scales — such as in cytoskeletal dynamics (3) — to tissular scales — such as in organ expansion (4). For instance, cell growth was found to be spatially heterogeneous (5-9), cell cycle length may appear random (10), and there is extensive evidence of stochastic gene expression (11, 12). Such variability has been hypothesised to be required for the emergence of complex shapes since it favors symmetry breaking (13) and self-organisation (14) during development. Nevertheless, growth variability would need to be restrained to ensure robust morphogenesis. In plant tissues, an increase in the spatial correlations of growth fluctuations was shown to reduce the robustness of floral organ size and shape (15). In animal tissues, work on the wing imaginal disc of the fruit fly indicates that robust wing development involves cell competition and requires the modulation of cell division and apoptosis (16, 17).

Mechanical signals are natural candidates for the regulation of growth variability because spatial differences in growth or in deformation rates induce mechanical stress (18-20). In animals, a mechanical feedback affecting the rate of cell divisions was hypothesized (21) and then supported by experiments in Drosophila and in zebrafish (22-25). Actomyosin cables are reinforced by mechanical tension in the wing imaginal disk of Drosophila (26). In plants, mechanical sensing is required to prevent growth fluctuations in roots (27). The deposition of cellulose fibers, the main load-bearing component of the cell wall, depends on wall tension (28, 29), which stiffens the cell wall in the direction of maximal tensile stress (30).

Previous theoretical studies have modelled how mechanical feedback regulates proliferation (21) and how transitions in tissue rheology are induced by proliferation and apoptosis (31, 32). Here, we build upon such studies; in addition, we account for small sources of stochasticity and investigate the consequences on large scale tissue growth. We focus on generic aspects of tissue growth, so that our results may be broadly applicable to active matter (33).

## GROWING TISSUES AS MECHANORESPONSIVE ACTIVE FLUIDS

We built a continuous two-dimensional model of tissue growth. The tissue is assumed to be made of a material with a preferred orientation (i.e. fiber-like), accounting for its main mechanical elements: cytoskeleton, polar cell-cell adhesive junctions, extra-cellular matrix (ECM) in animals; cellulose within the cell wall in plants. Hence, the state of the tissue is locally described by two order parameters, the density of fibers and the nematic field describing the orientation of fibers and their degree of alignment, which confer isotropic and anisotropic mechanical properties to the material, respectively. In our continuous description, we only account for variations in density, orientation, and degree of alignment at supra-cellular length scales, though the material may be patterned at smaller scales (sub-cellular or cellular). We account for fiber synthesis and remodeling, which may be modulated by responses to mechanical stress: reinforcement of actin stress fibers or of the ECM, enhancement of myosin activity, or fluidisation by cell division, in animals; increase in cell wall synthesis, cellulose synthesis, or cell division, in plants. Synthesis has a small random component, considered as a stochastic, uncorrelated source. Stochasticity in synthesis induces growth heterogeneity, which results in mechanical stress and feeds back on synthesis. We use a viscous description of long-term tissue remodeling, so that we cannot account for short-term elastic tissue behavior, which would be better captured by an elastic model as performed in a parallel study (34). Formally, the model describes an expanding, mechanoresponsive, nematic, and active fluid.

### A model of nematic viscous fluid

We describe the fibers with a density 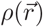 and a nematic 2 × 2 tensor 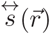, that vary with the position vector 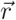. The nematic tensor 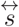 may be defined as an average over a small region around position 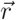, 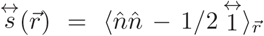, where the unit vector 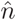 defines the polarization of fiber monomers and 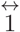 is the unit tensor. Regions of the material move at velocity 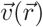, which may also vary spatially. In the following, we use the gradient of the velocity field, decomposed into strain rate, 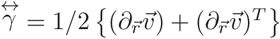, and vorticity 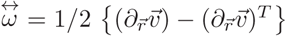, where 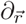 stands for the partial derivative with respect to position 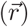 and *^T^* for the transpose of the preceding tensor.

We neglect diffusion of fibers in the tissue. The equations of continuity for density and nematic tensor are then

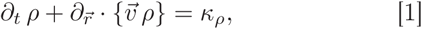

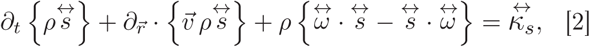

where *t* is time, *κ_ρ_* is the rate of synthesis of material, and 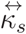 is a nematic tensor that describes the orientation of synthesis and its degree of alignment. The second terms in the left-hand sides of [1-2] account for the dilution of material density and degree of alignment due to tissue expansion; the third term of [2] accounts for the rotation of fibers due to the flow.

Expansion of the tissue is assumed to be driven by a uniform and isotropic tension, *p*, which may correspond to turgor pressure in plants, or to a pressure induced by cell divisions in animals (31); this tension is one of the active components of our model. The mechanical stress, 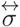, then follows the force balance equation 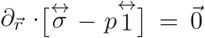, supplemented with standard boundary conditions: no shear stress and normal stress equal to tension, *p*, at system boundaries. We consider time scales long enough for tissue remodeling to occur, so that we neglect elastic behavior, assuming that 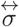 depends on the strain rate tensor, on the density, and on the nematic tensor. This dependence 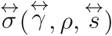 is the constitutive law that characterizes the rheology of the tissue.

In the following, we consider small fluctuations around an average state. The statistical averages of variables are denoted by brackets. For convenience, tensorial fields 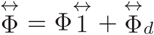 are decomposed into hydrostatic 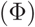 and deviatoric 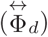 components, 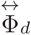 being traceless. On average, the tissue has uniform density, 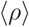, and is isotropic, 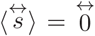; mechanical stress 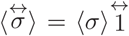 has only a hydrostatic component 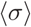; areal growth rate 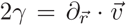 is on average uniform; through an appropriate change of reference frame, the averaged velocity may be written as 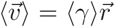. Assuming small fluctuations of all fields, we linearize the constitutive equation as a function of the hydrostatic strain rate, γ, the deviatoric strain rate, 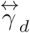, the density, ρ, and the nematic tensor 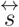,

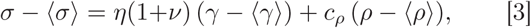

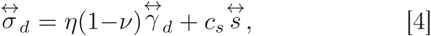

where *η* is an effective viscosity coefficient, *c_ρ_* and *c_s_* are effective compressibilities, and *v* is analog to Poisson’s ratio. In the right-hand side of [3-4], the first term accounts for viscous-like remodelling and the second for tissue compressibility (changes in density and alignment under stress).

### Activity: mechanical responses and fluctuations

On the one hand, mechanical stress orients cell divisions (22, 23, 35) and plant cell wall reinforcement (30). On the other hand, synthesis of ECM or of cell wall and cytoskeleton polymerization are not uniform in space, having some level of randomness (3, 36, 37). The two classes of phenomena are incorporated in the other active component of our model, namely synthesis. Without loss of generality, synthesis may be written at linear order in fluctuations as

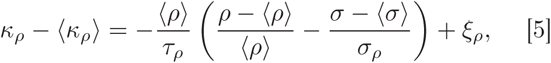

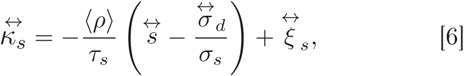

where the first terms in the right-hand side of [5-6] describe the mechanical feedback on synthesis. *τ_ρ_* and *τ_s_* are the response times of the mechanical feedbacks, *σ_ρ_* and *σ_s_* determine the amplitudes of the mechanical feedbacks, and *ξ_ρ_* and 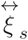 are the hydrostatic and deviatoric part of the noise, respectively. Noise is assumed to be white and Gaussian with zero mean, and has extended space correlation characterized by noise strengths *K_ρρ_* and *K_ss_* and by a correlation length *ℓ*, which is typically sub-cellular or cellular. The correlations functions of noise take the form 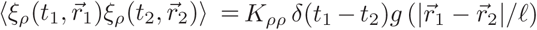, and 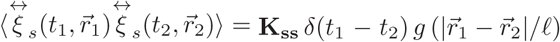. *δ* is the delta distribution and *g*(*x*) is a positive function decaying quickly to zero as *x* → +∞. (We use *g*(*x*) = *e*^−^*^x^* in calculations.) Cartesian coordinates of the 4-tensor **K_ss_** are constrained by the traceless nature of 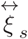 to be of the form **K_SS_***_abcd_* = *K_ss_*{*δ_ad_δ_bc_* + *δ_ac_δ_bd_* − *δ_db_δ_cd_*}, where *δ_ij_* is the Kronecker delta. We do not take the limit *ℓ* → 0 for spatial correlations because otherwise the problem would have no characteristic lengthscale. Accordingly, we hereafter use *ℓ* as a unit of length.

### Dimensionless parameters

We rescale all fields and variables as follows.

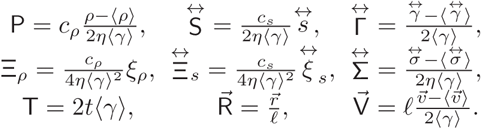

P and 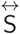 are the dimensionless density fluctuation and nematic tensor. They are associated to the dimensionless random components of synthesis 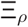 and 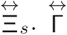 is the dimensionless fluctuation of strain rate tensor, 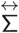 is the dimensionless stress fluctuation. T, 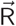, and 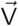 are, respectively, the dimensionless time, position vector, and velocity fluctuation. The dimensionless versions of Eqs. [1-6] are given in **[SI]**.

This rescaling shows that the model has 8 dimensionless parameters. *ω_ρ_* = 1 + 1/(2*τ_ρ_*〈*γ*〉) and *ω_s_* = 1 + 1/(2*τ_s_*〈*γ*〉) characterize the relaxation of the tissue in absence of mechanical feedback. *β*_0_ = *c_ρ_*〈*ρ*〉/(2*η*〈γ〉) compares the contributions of density variations and growth to mechanical stress. *v* is the dimensionless difference between effective dilatational and shear viscosities. K*_ρρ_* = *Kρρ*/(16*η*^2^〈*γ*〉^4^) and K*ss* = *Kss*/(16*η*^2^〈*γ*〉^4^) are the rescaled magnitudes of random synthesis. *β_ρ_* = *c_ρ_*〈*ρ*〉/(2*τ_ρ_*〈*γ*〉*σ_ρ_*) and *β_s_* = *c_s_*/(2*τ_s_*〈*γ*〉*σ_s_*) are the measures of isotropic and anisotropic responses to stress.

## RESPONSE TO PERTURBATIONS IN SYNTHESIS

The general formulation is given in the Appendix. Here, we discuss tissue response to an isotropic perturbation that is localised in space – a disk of initial radius *ℓ* – and in time – a duration that is small with respect to all other time scales. Formally, the perturbation to density synthesis is 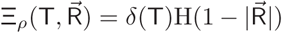, with H the Heaviside function, while the perturbation to synthesis of nematic order vanishes, 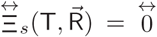. The fields 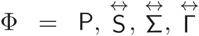 have self-similar forms, 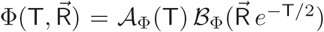, where 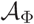 represents the amplitude of the perturbation and 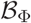 its spatial patterns. The dynamics of the amplitude is specific to each field, whereas the pattern always expands with a characteristic lengthscale *ℓ* exp(〈*γ*〉*t*) (in dimensional units). 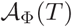 and 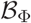 are represented in Fig. 1a-e and are explicitly given in **[SI]**. An immediate consequence of the perturbation is to stiffen the tissue, which reduces expansion 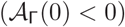 and increases stress levels 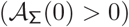; then the anisotropic mechanical response gradually induces radial fibers and reinforcement in the direction of the main stress. The behavior at longer times depends on the level of mechanical response. For low anisotropic response, i.e. for *β_s_* smaller than a threshold computed in **[SI]**, the tissue is hypo-responsive and all amplitudes evolve monotonously as a function of time and vanish at times that are long with respect to the correlation time *τ_c_*; tissue nematic orientation, strain rate, and mechanical stress are all mainly radial. For high anisotropic response, i.e. for *β_s_* above this threshold, the tissue is hyper-responsive, and amplitudes show an underdamped-like dynamics: they change sign before decaying to 0; after well-defined times, density becomes slightly smaller than average density, and all of nematic order, strain rate, and mechanical stress become circumferential. This hyper-responsive regime can be understood as follows. An initially high mechanical anisotropy of the tissue reduces the radial strain until strain becomes circumferential, leading to circumferential stress and then circumferential nematic order.

**FIG. 1.**
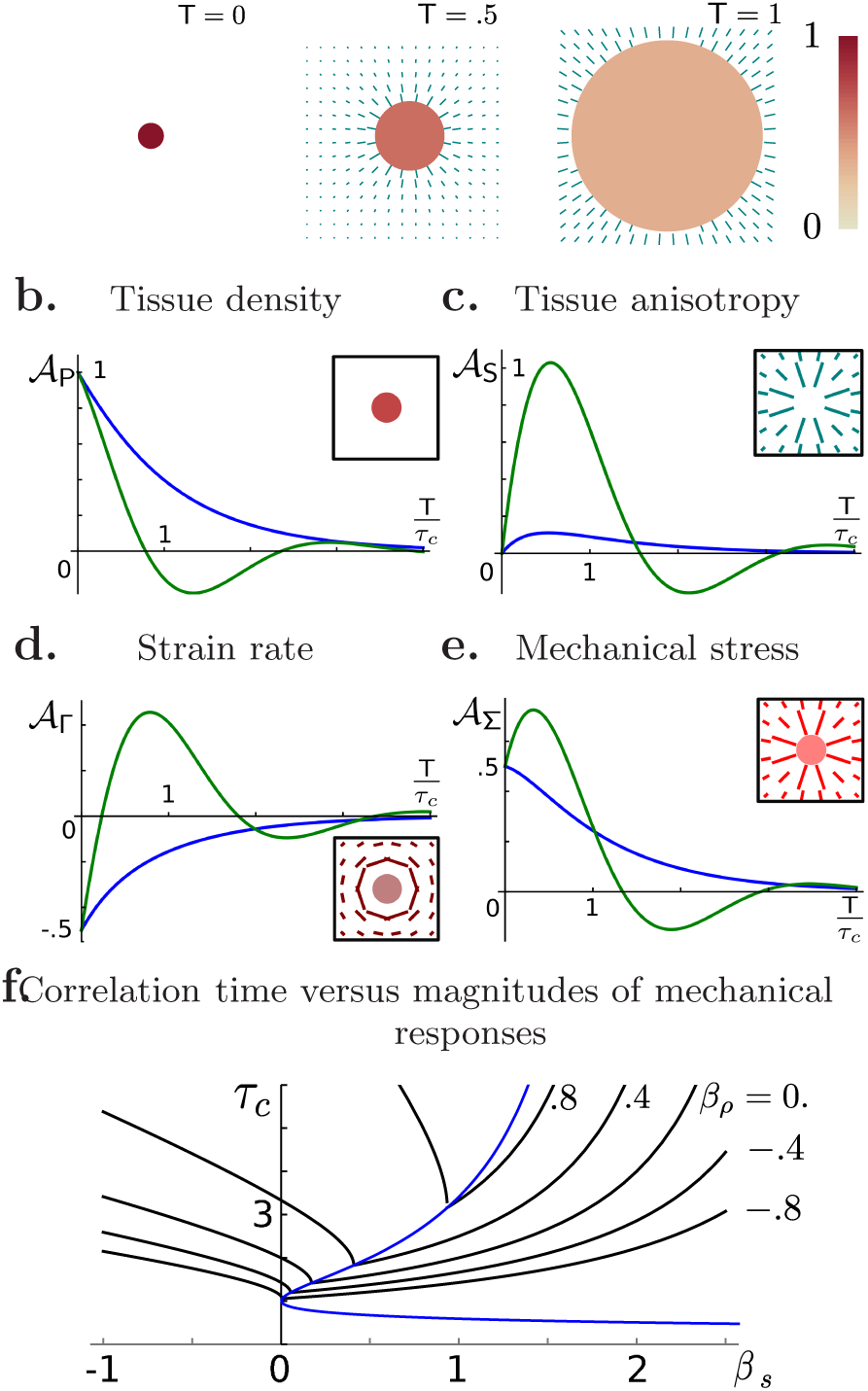
Example of mechanical response: tissue relaxation following a localized disk-shape isotropic perturbation, **a**. Snapshots at dimensionless times *T =* 0, 0.5 and 1. The density is color-coded according to the heat map on the left (white corresponds to no deviation from average density). The nematic order parameter is shown by the small lines: the angle corresponds to line orientation and the degree of anisotropy to line length. The mechanical responses strengths are *β_s_* = .3 and *β_ρ_* = .6. **b-e** Amplitudes 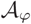 of the perturbations in tissue density, P, tissue anisotropy, S, strain rate, Γ, and mechanical stress, Σ; the corresponding patterns 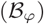 are shown as insets. The blue and the green lines show the relaxation of the amplitude of the perturbations for low, *β_s_* = 0.3, and high, *β_s_* = 1.6, anisotropic response, with *β_ρ_* = .6. Time is rescaled by the characteristic time, *τ_c_*. **f**. The correlation time, *τ_c_*, as a function of the strength of the isotropic (*β_ρ_*) and anisotropic (*β_s_*) mechanical responses. Regimes of hypo- and hyper-response are respectively on the left and on the right of the dashed blue line. *ω_ρ_* = 1, *ω_s_* = 1, *β*_0_ = 1, and *v* = 0 for all panels.

These dynamics occur on a time scale *τ_c_*, which is the maximal relaxation time scale in response to a perturbation **[SI]**. The time scale *τ_c_* depends on the magnitudes of mechanical responses, as shown in Fig. 1f (see Fig. S2 for the effect of other parameters). Isotropic mechanical feedback makes perturbations more persistent in time, because *τ_c_* increases with *β_ρ_*. The effect of the anisotropic feedback on relaxation is more complex: *τ_c_* first decrease and then increase as *β_s_* is increased; the minimum of *τ_c_* corresponds to the transition between hypo-response and hyper-response. This characteristic time *τ_c_* will also appear to be important for the effect of noise.

## GROWTH FLUCTUATIONS

We find that growth (areal strain rate in 2D) has long-range correlations, with a correlation function that spatially decays with an exponent −4/*τ_c_* **[SI]**. In practice, growth is measured at the scale of the spatial resolution of experimental measurements, which depends on the landmarks used and is often at cell scale. We therefore define a coarse-grained growth rate and we consider the time correlation function *G*(R, T) of the growth of a disk of radius R, where R is the coarse-graining size i.e. the resolution size. It is simply related to velocity fluctuations (see Appendix) by 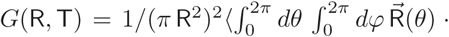. 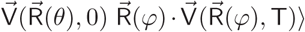. This correlation function is plotted in Fig. 2. Panel **a** shows time correlations for high and low anisotropic mechanical feedback, respectively corresponding to the hypo-response and hyper-response. The negative correlations for high feedback are related to underdamped relaxation of hyper-responsive tissues. The correlation function decays quickly to 0 with a characteristic time scale that is exactly the relaxation time, *τ_c_*, shown in Fig. 2e. Areal growth mean square deviation appears roughly scale-invariant, see Fig. 2b; it is exactly scale-invariant for hypo-response and oscillates around a scale-invariant for hyper-response. Two regimes characterise the decay. In a weakly-correlated regime, when *τ_c_* < 2, growth mean square deviation scales with the inverse of the coarse-graining area, *G*(R,0) ~ R^−2^, an exponent due to the central limit theorem. In a strongly-correlated regime, when *τ_c_* > 2, growth mean square deviation decays more slowly 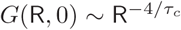, see **[SI]** for a rationale.

**FIG. 2.**
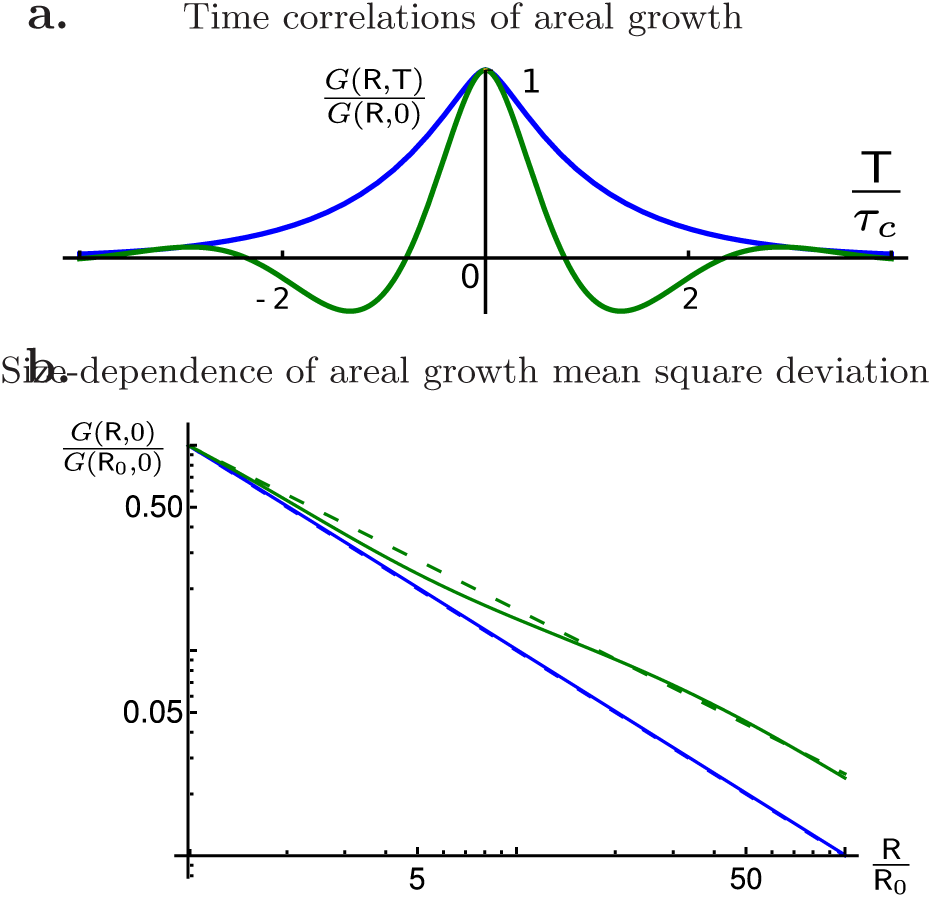
Growth fluctuations for low (*β_s_* = 0.3, blue) and high (*β_s_* = 1.6, green) anisotropic mechanical response. **a**. Time correlation function *G*(R,T) (normalized by its initial value) as a function of time, T, normalized by the correlation time, *τ_c_*; see Fig. S1 for other values of *β_s_*. Negative correlations appear for high anisotropic mechanical response. **b**. Growth mean square deviation, *G*(R,T), as a function of the coarse-graining size, R. *G*(R,T) and R are normalized using R_0_ = 20. The asymptotic power-law (R^−1.2^) for low anisotropic mechanical response is shown by the blue dashed line. For high anisotropic mechanical response, *G*(R,T) oscillates around a power-law (R^−0.8^, dashed green line). *β_ρ_* = 0.6, *ω_ρ_* = 1, *ω_s_* = 1, *β*_0_ = 1, and *ν* = 0 for the two panels.

## FLUCTUATIONS OF ORGAN SHAPE

We are now interested in the effects of noise in synthesis on tissue contours or on organ shape. In a homogeneous and isotropic tissue, quantifying the fluctuation of contours is equivalent to determining the fluctuation of a vector joining two landmarks followed throughout growth of the tissue. Hence, we use a Lagrangian description and consider the position, 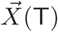, at time *T* of a landmark initially at position 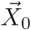, which is determined by the dimensionless velocity field 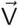 through 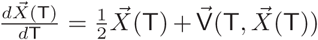. The fluctuations of 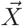 are computed in the Appendix. Heterogeneity of contours is assessed using the coefficient of variation, 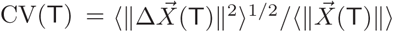, which is plotted in Fig. 3a. Its asymptotic trend for long times depends on the value of the correlation time *τ_c_*. If *τ_c_* < 2, then 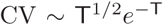, and for *τ_c_* > 2, CV scales 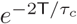 for low feedback and oscillate around this trend for high feedback.

**FIG. 3.**
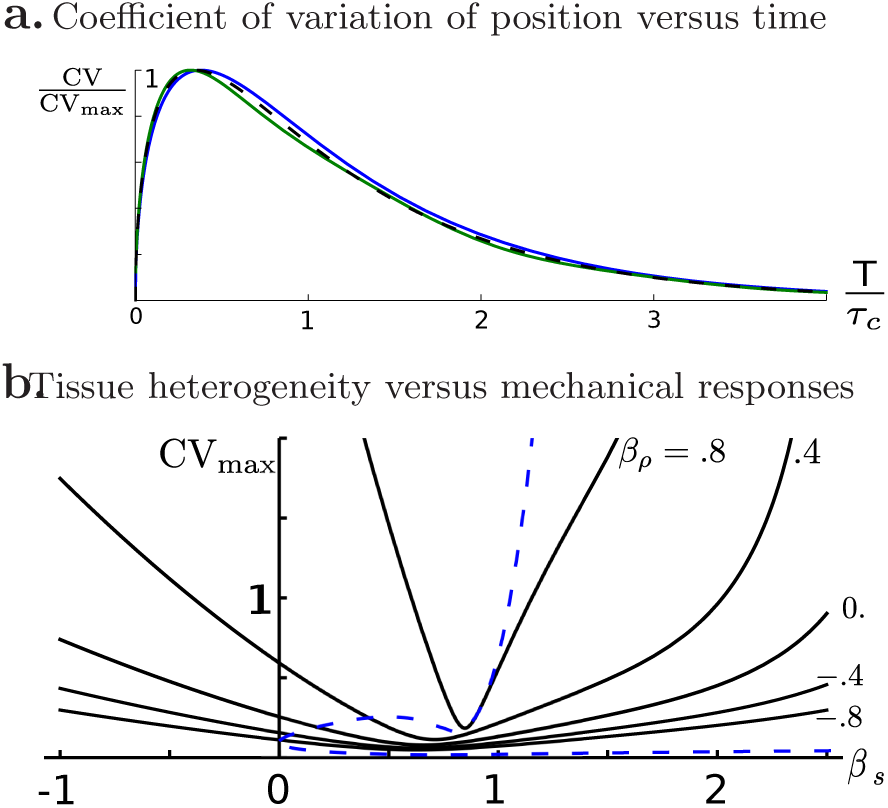
**a**. Coefficient of variation of position, 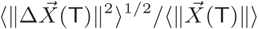, normalized by its maximal value CV_max_ as a function of time, T, normalized by the correlation time, *τ_c_*, for low (*β_s_* = 0.3, blue) and high (*β_s_* = 1.6, green) anisotropic mechanical response (see Fig. S3 for other values of *β_s_*). The dashed line represents the asymptotic limit for *X*_0_ ≫ 1 for low anisotropic feedback. **b**. Coefficient of variation of position, normalized by 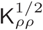, as a function of the magnitude of anisotropic mechanical response for various levels of anisotropic feedback. Regimes of hypo- and hyper-response are respectively on the left and on the right of the dashed blue line. *β_ρ_* = 0.6, *ω_ρ_* = 1, *ω_s_* = 1, *β*_0_ = 1, *ν* = 0, and *X*_0_ = 10 for the two panels.

We represent in Fig. 3b the maximal value of the coefficient of variation, normalized by 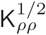. This enables us to quantify contour fluctuations in the tissue or of organ shapes. In absence of anisotropic feedback, *β_s_* = 0, we find that heterogeneity increases with isotropic feedback. Accordingly, a positive isotropic feedback maintains perturbations and induces long-range correlations as seen in previous sections. Conversely, negative isotropic feedback dampens perturbations. Whatever the level of isotropic feedback, *β_ρ_*, we find that heterogeneity, as a function of anisotropic feedback, has a single minimum, which corresponds to the transition between hypo- and hyper-response. In the hypo-responsive regime, increasing anisotropic feedback dampens perturbations, whereas in the hyper-responsive regime, increasing anisotropic feedback enhances perturbations due to the oscillatory overshoot. Finally, we note that the behavior of heterogeneity in Fig. 3b is qualitatively similar to the behavior of correlation time (*τ_c_*) in Fig. 1f, indicating that the correlation time is a major determinant of heterogeneity because the correlation time sets how the tissue keeps the memory of its previous state.

## DISCUSSION

We built a continuous viscous model of tissue growth, describing density and nematic order of the tissue, and modelled material synthesis and responses to mechanical stress. Here, the responses are characterized by two parameters, *β_ρ_* and *β_s_*, corresponding to isotropic response - increase in density due to increase in stress when *β_ρ_* > 0 - and anisotropic response - increase in tissue anisotropy due to increase in stress anisotropy when *β_s_* > 0, and conversely when these parameters are negative. In plants, it is believed that cell wall synthesis is enhanced when tension increases (38), which corresponds to *β_ρ_* > 0. The alignment of cortical microtubules with maximal stress orientation leads to the anisotropic stiffening of the cell wall in this direction (30, 39), while cell divisions are associated with new cell walls oriented in the direction of maximal stress (35); both processes yield *β_s_* > 0. In animal tissues, experiments indicate that tissues are fluidised by cell divisions (22-25): proliferation is enhanced by tensile stress and daughter cells tend to separate along the direction of highest mechanical stress, which corresponds to *β_ρ_* < 0 and *β_s_* < 0, respectively. At shorter time scales, actomyosin cables are reinforced in the direction of applied stress (26), which yields *β_ρ_* > 0. At intermediate time scales, ECM would also be reinforced against mechanical stress (*β_ρ_* > 0), though its role in morphogenesis has not received attention until recently (40-43).

In this study, we determined tissue response to a localized perturbation, depending on the mechanical feedback parameters *β_ρ_* and *β_s_*. We generalized predictions that anisotropic mechanical feedback buffers such a perturbation (29), in agreement with observations on trichomes – a cell type with transient faster growth -–in Arabidopsis sepals (29). Here, we unravelled two possible regimes: hypo-response at low anisotropic feedback – perturbations decay monotonously and hyper-response at high anisotropic feedback – perturbations oscillate before decaying, with a characteristic time that is minimal at the transition between the two regimes. This case study provides an assay of mechanical responses in both plant and animal systems, for instance by inducing clones with altered growth rate and quantifying the relaxation timescales in backgrounds with different levels of mechanical response.

We then investigated the statistical properties of tissue growth, unravelling long-range correlations, with slowly decaying correlation functions. To test this, it would be crucial to examine correlation functions in live imaging data of growing organs, e.g. (15, 44-47). Given that a larger correlation time (*τ_c_*) corresponds to longer-range correlations, higher levels of anisotropic feedback (independently of its sign) or higher levels of positive isotropic feedback yield more slowly decaying correlations functions. Nevertheless, long-range correlations could be mediated by both chemical and mechanical signals. Further experiments would be required to test whether mechanical signals are involved in such correlations.

Finally, we found that heterogeneity of contours and shapes is minimal for a well-determined level of anisotropic mechanical response. This generalizes a similar conclusion reached for local heterogeneity using a cell-based toy model (48). Here we also accounted for isotropic mechanical responses and considered heterogeneity at all scales. We identified the correlation time as a key parameter determining the extent of spatial correlations and the level of heterogeneity of organ shape. Based on our results, we make the following predictions. In plants (*β_ρ_* > 0 and *β_s_* > 0), heterogeneity in development can be significantly high unless anisotropic feedback is close to the value that minimizes heterogeneity. In animals, if we discard possible contributions of the ECM, *β_ρ_* < 0 and *β_s_* < 0 at long time scales, heterogeneity in development is minimal when anisotropic feedback is negligible. Such predictions can be tested by measuring correlations of growth in space and time, as well as the strength of isotropic and anisotropic variations in growth due to an external force or to the induction of clones. More generally, characterizing the fluctuations of cell properties appears as a promising avenue to shed light on how signals orchestrate organismal development.

## Supporting information

## APPENDIX

### Response to perturbations in synthesis

We assume the tissue to be infinite, the perturbations to not induce rotation 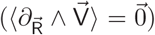, and the reference frame to satisfy 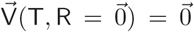. We investigate the response to generic perturbations. Because the average strain rate profile stretches patterns, we consider a modified Fourier transform defined as

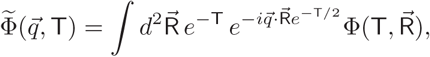

with the position 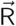 rescaled by the average growth factor e^T/2^. Defining 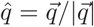 as the direction of the wavevector, the Fourier transform of the nematic tensor can be written as 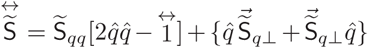. The linear response for material density and nematic tensor is then given by

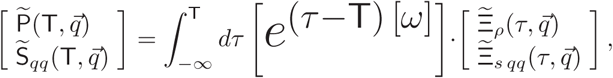

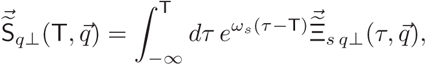

where 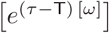 is an exponential involving the relaxation matrix 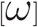, see **[SI]**, and the fields 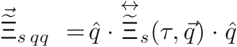 and 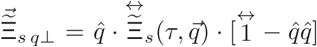 are the components of the noise Fourier transform. Finally, the Fourier transform of the strain rate tensor is decomposed as 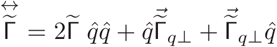, so that the linear response of strain rate is given by

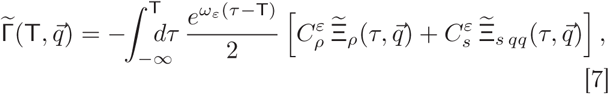

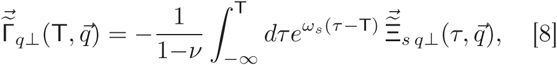

where the integrand in the r.h.s. of [7] is a sum over the values {+, −} of the index *ε* and *ω_±_* are the two eigenvalues of [*ω*]. The expression of *ω_±_* and the coefficients 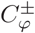 can be found in **[SI]**. We thus obtain the full response of the tissue to any perturbation of synthesis, in terms of the modified Fourier transform of the sources of density and of nematic order.

#### Growth fluctuations

Using the linear response of flow velocity to synthesis perturbations [7-8], we derived the velocity fluctuations, as detailed in **[SI]**. The correlation tensor of velocity fluctuations is proportional to the unit tensor,

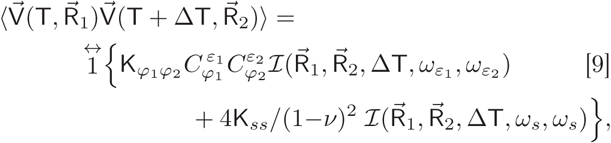

for ΔT > 0, with the same 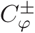 coefficients as in [7-8] and

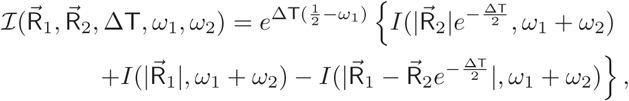

where 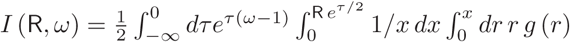. This explicitly defines the correlation functions.

#### Fluctuations of organ shape

We look for the probability 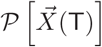 that a material point follows a path 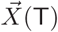. For small fluctuations, this probability can be simplified as

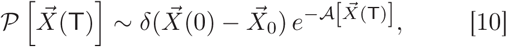

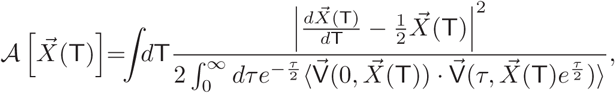

where the velocity correlation function is given by [9]. We determined the asymptotic statistics of the Lagrangian flow by applying the saddle point method (49) to 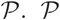 is maximized by the average trajectory 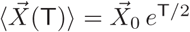 and the correlation tensor of the position 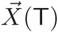 is given by [SI]

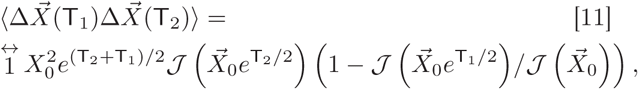

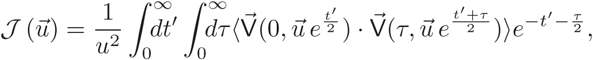

with T_1_ ≤ T_2_. This correlation tensor does not depend on T_2_ − T_1_ because of the lack of time translation invariance.

## References

[1] H. M. Meyer and A. H. Roeder, Frontiers in plant science 5, 420 (2014).

[2] A. C. Oates, Development 138, 601 (2011).

[3] D. S. Banerjee, A. Munjal, T. Lecuit, and M. Rao, Nature Communications 8, 1121 (2017).

[4] M. Sahaf and E. Sharon, Journal of experimental botany 67, 5509 (2016).

[5] M. Uyttewaal, A. Burian, K. Alim, B. Landrein, D. Borowska-Wykręt, A. Dedieu, A. Peaucelle, M. Ludynia, J. Traas, A. Boudaoud, et al., Cell 149, 439 (2012).

[6] G. Tauriello, H. M. Meyer, R. S. Smith, P. Koumoutsakos, and A. H. Roeder, Plant physiology 169, 2342 (2015).

[7] D. Kierzkowski, N. Nakayama, A.-L. Routier-Kierzkowska, A. Weber, E. Bayer, M. Schorderet, D. Reinhardt, C. Kuhlemeier, and R. S. Smith, Science 335, 1096 (2012).

[8] J. Elsner, M. Michalski, and D. Kwiatkowska, Annals of botany 109, 897 (2012).

[9] S. L. Martin, Nature 460, 1087 (2009).

[10] A. H. Roeder, V. Chickarmane, A. Cunha, B. Obara, B. Manjunath, and E. M. Meyerowitz, PLoS biology 8, e1000367 (2010).

[11] A. Raj and A. van Oudenaarden, Cell 135, 216 (2008).

[12] I. S. Araújo, J. M. Pietsch, E. M. Keizer, B. Greese, R. Balkunde, C. Fleck, and M. Hülskamp, Nature communications 8, 2132 (2017).

[13] F. Corson, L. Couturier, H. Rouault, K. Mazouni, and F. Schweisguth, Science 356, eaai7407 (2017).

[14] J. R. Chubb, Wiley Interdisciplinary Reviews: Developmental Biology 6 (2017).

[15] L. Hong, M. Dumond, S. Tsugawa, A. Sapala, A.-L. Routier-Kierzkowska, Y. Zhou, C. Chen, A. Kiss, M. Zhu, O. Hamant, et al., Developmental cell 38, 15 (2016).

[16] D. R. Hipfner and S. M. Cohen, Nature Reviews Molecular Cell Biology 5, 805 (2004).

[17] M. Milán, S. Campuzano, and A. García-Bellido, Proceedings of the National Academy of Sciences 93, 640 (1996).

[18] K. D. Irvine and B. I. Shraiman, Development 144, 4238 (2017).

[19] V. Mirabet, P. Das, A. Boudaoud, and O. Hamant, Annual review of plant biology 62, 365 (2011).

[20] G.-J. J. Gao, M. C. Holcomb, J. H. Thomas, and J. Blawzdziewicz, Journal of Physics: Condensed Matter 28, 414021 (2016).

[21] B. I. Shraiman, Proceedings of the National Academy of Sciences of the United States of America 102, 3318 (2005).

[22] L. LeGoff, H. Rouault, and T. Lecuit, Development 140, 4051 (2013).

[23] P. Campinho, M. Behrndt, J. Ranft, T. Risler, N. Minc, and C.-P. Heisenberg, Nature cell biology 15, 1405 (2013).

[24] Y. Mao, A. L. Tournier, A. Hoppe, L. Kester, B. J. Thompson, and N. Tapon, The EMBO journal 32, 2790 (2013).

[25] Y. Pan, I. Heemskerk, C. Ibar, B. I. Shraiman, and K. D. Irvine, Proceedings of the National Academy of Sciences 113, E6974 (2016).

[26] M. Duda, N. Khalilgharibi, N. Carpi, A. Bove, M. Piel, G. Charras, B. Baum, and Y. Mao, bioRxiv, 241497 (2017).

[27] H.-W. Shih, N. D. Miller, C. Dai, E. P. Spalding, and G. B. Monshausen, Current Biology 24, 1887 (2014).

[28] O. Hamant, M. G. Heisler, H. Jönsson, P. Krupinski, M. Uyttewaal, P. Bokov, F. Corson, P. Sahlin, A. Boudaoud, E. M. Meyerowitz, et al., science 322, 1650 (2008).

[29] N. Hervieux, S. Tsugawa, A. Fruleux, M. Dumond, A.-L. Routier-Kierzkowska, T. Komatsuzaki, A. Boudaoud, J. C. Larkin, R. S. Smith, C.-B. Li, et al., Current Biology 27, 3468 (2017).

[30] A. Sampathkumar, P. Krupinski, R. Wightman, P. Milani, A. Berquand, A. Boudaoud, O. Hamant, H. Jönsson, and E. M. Meyerowitz, eLife 3, e01967 (2014).

[31] J. Ranft, M. Basan, J. Elgeti, J.-F. Joanny, J. Prost, and F. Jülicher, Proceedings of the National Academy of Sciences 107, 20863 (2010).

[32] D. Matoz-Fernandez, E. Agoritsas, J.-L. Barrat, E. Bertin, and K. Martens, Physical review letters 118, 158105 (2017).

[33] M. C. Marchetti, J.-F. Joanny, S. Ramaswamy, T. B. Liverpool, J. Prost, M. Rao, and R. A. Simha, Reviews of Modern Physics 85, 1143 (2013).

[34] O. Khatib Damavandi and D. K. Lubensky, bioRxiv (2018).

[35] M. Louveaux, J.-D. Julien, V. Mirabet, A. Boudaoud, and O. Hamant, Proceedings of the National Academy of Sciences 113, E4294 (2016).

[36] B. Altartouri and A. Geitmann, Current opinion in plant biology 23, 76 (2015).

[37] C. T. Anderson, I. S. Wallace, and C. R. Somerville, Proceedings of the National Academy of Sciences 109, 1329 (2012).

[38] S. Wolf, K. Hématy, and H. Höfte, Annual review of plant biology 63, 381 (2012).

[39] B. Landrein and O. Hamant, The Plant Journal 75, 324 (2013).

[40] J. Crest, A. Diz-Muñoz, D.-Y. Chen, D. A. Fletcher, and D. Bilder, Elife 6, e24958 (2017).

[41] R. P. Ray, P. S. Ganguly, S. Alt, J. R. Davis, A. Hoppe, N. Tapon, G. Salbreux, B. J. Thompson, et al., Developmental cell 46, 23 (2018).

[42] J. Chlasta, P. Milani, G. Runel, J.-L. Duteyrat, L. Arias, L.-A. Lamiré, A. Boudaoud, and M. Grammont, Development, dev (2017).

[43] R. Loganathan, B. J. Rongish, C. M. Smith, M. B. Filla, A. Czirok, B. Bénazéraf, and C. D. Little, Development 143, 2056 (2016).

[44] I. Heemskerk, T. Lecuit, and L. LeGoff, Development 141, 2339 (2014).

[45] B. Guirao, S. U. Rigaud, F. Bosveld, A. Bailles, J. López-Gay, S. Ishihara, K. Sugimura, F. Graner, and Y. Bellaïche, Elife 4, e08519 (2015).

[46] E. Rozbicki, M. Chuai, A. I. Karjalainen, F. Song, H. M. Sang, R. Martin, H.-J. Knölker, M. P. MacDonald, and C. J. Weijer, Nature cell biology 17, 397 (2015).

[47] E. E. Kuchen, S. Fox, P. B. De Reuille, R. Kennaway, S. Bensmihen, J. Avondo, G. M. Calder, P. Southam, S. Robinson, A. Bangham, et al., Science 335, 1092 (2012).

[48] K. Alim, Frontiers in Plant Science 3, 1 (2012).

[49] A. Altland and B. D. Simons, Condensed matter field theory (Cambridge University Press, 2010) Chap. 3, pp. 108–111.

