## Supplementary material for "Modulation of tissue growth heterogeneity by responses to mechanical stress"

### Supplementary information for:
#### Modulation of tissue growth heterogeneity by responses to mechanical stress

Antoine Fruleux and Arezki Boudaoud*

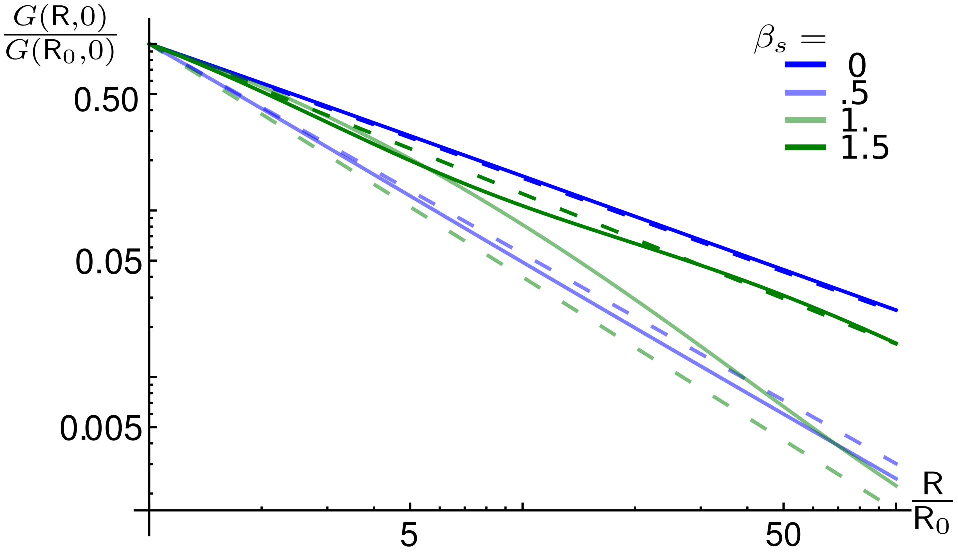


FIG. 1. Growth mean square deviation, $G(\mathsf{R},\mathsf{T})$, as a function of the coarse-graining size, $\mathsf{R}$, shown for different levels of anisotropic mechanical response, $\beta_s$. $G(\mathsf{R},\mathsf{T})$ and $\mathsf{R}$ are normalized using $\mathsf{R}_0 = 20$. Blue and green lines correspond to hypo- and hyper-responsive regimes, respectively. The asymptotic power-law for low anisotropic mechanical response is shown by the blue dashed line. For high anisotropic mechanical response, $G(\mathsf{R},\mathsf{T})$ oscillates around a power-law (dashed green lines). $\omega_\rho = 1$, $\omega_s = 1$, $\beta_0 = 1$, $\beta_\rho = .6$, and $\nu = 0$.

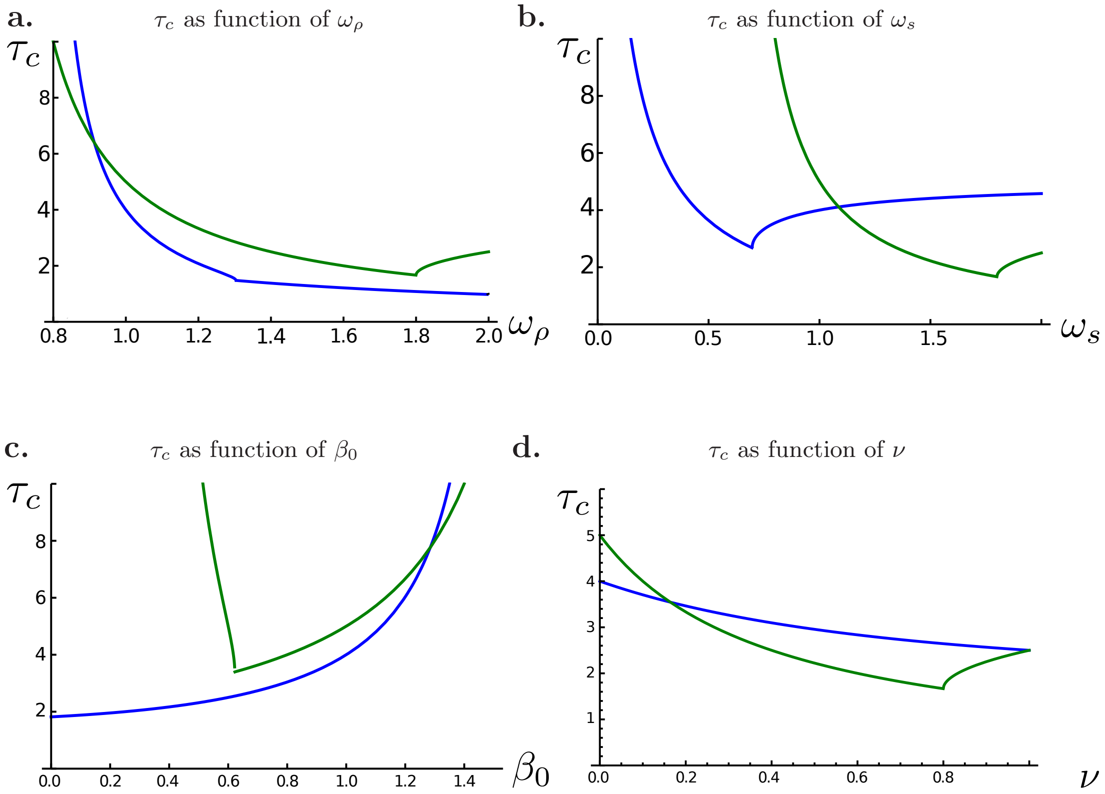


FIG. 2. The correlation time, $\tau_c$, as a function of model parameters: **a.** $\omega_\rho$; **b.** $\omega_s$; **c.** $\beta_0$; **d.** $\nu$. Blue and green lines correspond to hypo- and hyper-responsive regimes, respectively. The reference values of parameters are $\beta_\rho = .6$, $\beta_s = 0.3$ or $\beta_s = 1.6$, $\omega_\rho = 1$, $\omega_s = 1$, $\beta_0 = 1$, and $\nu = 0$, for all panels.

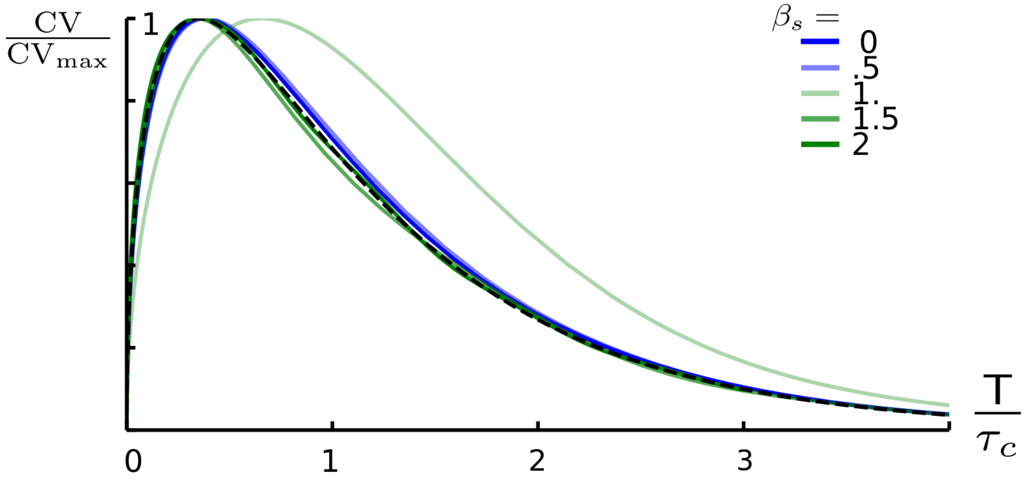


FIG. 3. Coefficient of variation of position, $\mathrm{CV} = \langle \|\Delta\vec{X}(\mathsf{T})\|^2\rangle^{1/2}/\langle\|\vec{X}(\mathsf{T})\|\rangle$, normalized by its maximal value $\mathrm{CV}_{\max}$ as a function of time, $\mathsf{T}$, normalized by the correlation time, $\tau_c$, shown for different levels of anisotropic mechanical response, $\beta_s$. Blue and green lines correspond to hypo- and hyper-responsive regimes, respectively. The dashed line represents the asymptotic limit for $X_0 \gg 1$ for low anisotropic feedback; it gives the good trend for most parameter values, though CV tend to deviate from this trend at the transition from hypo- to hyper-response.

This supplementary text provides details of models and derivations presented in the main text. The titles of sections and subsections generally follow the titles of the main text.

#### GROWING TISSUES AS MECHANORESPONSIVE ACTIVE FLUIDS

##### A model of nematic viscous fluid

We describe two-dimensional tissues with a material density $\rho$ and a material anisotropy represented with a nematic tensor $\overleftrightarrow{s}$. Each material at coordinates point has a velocity $\vec{v}$, so that density and nematic tensor are advected, leading to the continuity equations

$$\partial_t \rho + \partial_{\vec{r}} \cdot \{\vec{v}\,\rho\} = \kappa_\rho, \qquad [1]$$

$$\partial_t \left\{\rho \overleftrightarrow{s}\right\} + \partial_{\vec{r}} \cdot \left\{\vec{v}\,\rho \overleftrightarrow{s}\right\} + \rho \left\{\overleftrightarrow{\omega} \cdot \overleftrightarrow{s} - \overleftrightarrow{s} \cdot \overleftrightarrow{\omega}\right\} = \overleftrightarrow{\kappa}_s. \qquad [2]$$

Here and elsewhere, $t$ stands for time and $\vec{r}$ for the position vector in the plane of the tissue; $\partial_X$ denotes the partial derivative with respect to variable $X$. $\kappa_\rho$ is the rate of synthesis of new material and $\overleftrightarrow{\kappa}_s$ is the rate of synthesis of anisotropic material. The last term in the l.h.s. of [2] accounts for the corotation of material anisotropy with fluid particles, according to flow vorticity $\overleftrightarrow{\omega} = 1/2\left\{(\partial_{\vec{r}}\vec{v}) - (\partial_{\vec{r}}\vec{v})^T\right\}$, where $^T$ stands for tensor transpose.

The mechanical balance between hydrostatic tension $p$ and tissue tension $\overleftrightarrow{\sigma}$ can be written as

$$\partial_{\vec{r}} \cdot \left[\overleftrightarrow{\sigma} - p\overleftrightarrow{1}\right] = \vec{0}, \qquad [3]$$

where $\overleftrightarrow{1}$ is the unit tensor. This equation is supplemented with standard boundary conditions: no shear stress and normal stress equal to tension $p$ at system boundaries. Cell wall tension is assumed to depend on material density, $\rho$, on material anisotropy, $\overleftrightarrow{s}$, and on the strain rate $\overleftrightarrow{\gamma} = \frac{1}{2}\left\{(\partial_{\vec{r}}\vec{v}) + (\partial_{\vec{r}}\vec{v})^T\right\}$, through the constitutive equation

$$\overleftrightarrow{\sigma} = \sigma\left(\overleftrightarrow{\gamma}, \rho, \overleftrightarrow{s}\right). \qquad [4]$$

##### Activity: mechanical responses and fluctuations

In our model, activity of the material relies on synthesis of new material. We consider deviations from average synthesis and consequences on tissue growth. A bracketed quantity, such as in $\langle X\rangle$, stands for its average. The tissue is assumed isotropic: material anisotropy vanishes on average, $\langle\overleftrightarrow{s}\rangle = \overleftrightarrow{0}$, and tension is isotropic, $\langle\overleftrightarrow{\sigma}\rangle = \langle\sigma\rangle\overleftrightarrow{1}$. For convenience, each tensor $\overleftrightarrow{\Phi}$ is decomposed into isotropic and deviatoric components, $\Phi$ and $\overleftrightarrow{\Phi}_d$, respectively, such that $\overleftrightarrow{\Phi} = \Phi\overleftrightarrow{1} + \overleftrightarrow{\Phi}_d$ where $\overleftrightarrow{\Phi}_d$ is traceless. Surface growth rate $2\gamma = \partial_{\vec{r}} \cdot v$ is on average constant. In the appropriate reference frame (and without loss of generality), averaged velocity is then $\langle\vec{v}\rangle = \langle\gamma\rangle\vec{r}$.

We expand synthesis at linear order in deviations of synthesis and stress, which leads to the general expressions,

$$\kappa_\rho = \langle\kappa_\rho\rangle - \frac{\langle\rho\rangle}{\tau_\rho}\left(\frac{\rho - \langle\rho\rangle}{\langle\rho\rangle} - \frac{\sigma - \langle\sigma\rangle}{\sigma_\rho}\right) + \xi_\rho, \qquad [5]$$

$$\overleftrightarrow{\kappa}_s = -\frac{\langle\rho\rangle}{\tau_s}\left(\overleftrightarrow{s} - \frac{\overleftrightarrow{\sigma}_d}{\sigma_s}\right) + \overleftrightarrow{\xi}_s, \qquad [6]$$

where $\tau_\rho$ and $\tau_s$ are response times for mechanical responses; $\sigma_\rho$ and $\sigma_s$ are constant coefficients determining response amplitude. $\xi_\rho$ and $\overleftrightarrow{\xi}_s$ stand for the noise on synthesis. Noise is assumed to be Gaussian, white in time and with localised correlations in space; it is characterized by amplitudes $K_{\rho\rho}$ and $K_{ss}$ and by a correlation length $\ell$:

$$\langle\xi_\rho(t_1, \vec{r}_1)\xi_\rho(t_2, \vec{r}_2)\rangle = K_{\rho\rho}\,\delta(t_1 - t_2)\,g\left(\frac{|\vec{r}_1 - \vec{r}_2|}{\ell}\right), \qquad [7]$$

$$\langle \overleftrightarrow{\xi}_s(t_1,\vec{r}_1)\overleftrightarrow{\xi}_s(t_2,\vec{r}_2)\rangle = \mathbf{K_{ss}}\,\delta(t_1-t_2)\,g\left(\frac{|\vec{r}_1-\vec{r}_2|}{\ell}\right). \quad [8]$$

The 4th order tensor $\mathbf{K_{ss}}$ is constrained by the traceless nature of $\overleftrightarrow{\xi}_s$ to be of the form $\mathbf{K_{ss}}_{abcd} = K_{ss}\,\{\delta_{ad}\delta_{bc}+\delta_{ac}\delta_{bd}-\delta_{ab}\delta_{cd}\}$ where $\delta_{ij}=1$ if $i=j$ and $\delta_{ij}=0$ if $i\neq j$. $g(x)$ is a positive function, defined for $x>0$, that decays quickly to 0 as $x>1$. In explicit calculations, we hereafter use $g(x)=e^{-x}$.

We assume fluctuations in synthesis to be small and we linearize the system of equations [1-2] around average. The linearised isotropic and deviatoric components of the stress are

$$\sigma = \langle\sigma\rangle + \eta(1{+}\nu)\,(\gamma-\langle\gamma\rangle) + c_\rho\,(\rho-\langle\rho\rangle), \quad [9]$$

$$\overleftrightarrow{\sigma}_d = \eta(1{-}\nu)\overleftrightarrow{\gamma}_d + c_s\overleftrightarrow{s}, \quad [10]$$

respectively, where $\eta$ is an effective viscosity coefficient; $c_\rho$ and $c_s$ are constant coefficients that reflect the dependance of rheology on ; $\nu$ is a constant that plays a similar role as Poisson's ratio.

#### Dimensionless parameters

In order to non-dimensionalize the set of equations, we defined rescaled material density, material anisotropy, strain rate, isotropic noise, anisotropic noise, stress, time, position vector, and velocity, as follows.

$$\begin{array}{lll} \mathsf{P} = c_\rho\frac{\rho-\langle\rho\rangle}{2\eta\langle\gamma\rangle}, & \overleftrightarrow{\mathsf{S}} = \frac{c_s}{2\eta\langle\gamma\rangle}\overleftrightarrow{s}, & \overleftrightarrow{\Gamma} = \frac{\overleftrightarrow{\gamma}-\langle\overleftrightarrow{\gamma}\rangle}{2\langle\gamma\rangle}, \\ \Xi_\rho = \frac{c_\rho}{4\eta\langle\gamma\rangle^2}\xi_\rho, & \overleftrightarrow{\Xi}_s = \frac{c_s}{4\eta\langle\gamma\rangle^2}\overleftrightarrow{\xi}_s, & \overleftrightarrow{\Sigma} = \frac{\overleftrightarrow{\sigma}-\langle\overleftrightarrow{\sigma}\rangle}{2\eta\langle\gamma\rangle}, \\ \mathsf{T} = 2t\langle\gamma\rangle, & \vec{\mathsf{R}} = \frac{\vec{r}}{\ell}, & \vec{\mathsf{V}} = \ell\frac{\vec{v}-\langle\vec{v}\rangle}{2\langle\gamma\rangle}. \end{array}$$

The final model (see below) has 8 non-dimensional parameters ($\omega_\rho$, $\omega_s$, $\beta_0$, $\nu$, $\mathsf{K}_{\rho\rho}$, $\mathsf{K}_{ss}$, $\beta_\rho$ and $\beta_s$) that are defined as follows. $\omega_\rho = 1+\frac{1}{2\tau_\rho\langle\gamma\rangle}$ and $\omega_s = 1+\frac{1}{2\tau_s\langle\gamma\rangle}$ compare the dynamics of synthesis and tissue expansion. $\beta_0 = \frac{c_\rho\langle\rho\rangle}{2\eta\langle\gamma\rangle}$ compares how material density fluctuations and strain rate contribute to mechanical stress. $\nu$ compares effective compression viscosity and shear viscosity. $\mathsf{K}_{\rho\rho} = \frac{K_{\rho\rho}}{16(\eta+\eta_d)^2\langle\gamma\rangle^4}$ and $\mathsf{K}_{ss} = \frac{K_{ss}}{16(\eta+\eta_d)^2\langle\gamma\rangle^4}$ are the rescaled noise amplitudes. Finally, $\beta_\rho = \frac{1}{2\tau_\rho\langle\gamma\rangle}\frac{c_\rho\langle\rho\rangle}{\sigma_\rho}$ and $\beta_s = \frac{1}{2\tau_s\langle\gamma\rangle}\frac{c_s}{\sigma_s}$ are the rescaled strengths of mechanical responses.

#### Dimensionless equations

When appropriate, tensors are decomposed as the sum of isotropic and deviatoric parts, such as the dimensionless deviatoric stress $\overleftrightarrow{\Sigma} = \Sigma\overleftrightarrow{1} + \overleftrightarrow{\Sigma}_d$.

Momentum conservation takes the form

$$\partial_{\vec{\mathsf{R}}}\cdot\overleftrightarrow{\Sigma} = \vec{0}. \quad [11]$$

$\overleftrightarrow{\Sigma}$ depends on the strain rate

$$\overleftrightarrow{\Gamma} = \frac{1}{2}\left\{(\partial_{\vec{\mathsf{R}}}\vec{\mathsf{V}}) + (\partial_{\vec{\mathsf{R}}}\vec{\mathsf{V}})^T\right\}, \quad [12]$$

on material density $\mathsf{P}$, and on nematic tensor $\overleftrightarrow{\mathsf{S}}$. More precisely, the isotropic and deviatoric tension are

$$\Sigma = (1{+}\nu)\Gamma + \mathsf{P}, \quad [13]$$

$$\overleftrightarrow{\Sigma}_d = (1{-}\nu)\overleftrightarrow{\Gamma}_d + \overleftrightarrow{\mathsf{S}}. \quad [14]$$

The dynamic equations for density and nematic tensor account for relaxation to reference state, for mechanical responses, and for noise in synthesis,

$$\partial_{\mathsf{T}}\,\mathsf{P} + \frac{1}{2}(\vec{\mathsf{R}}\cdot\partial_{\vec{\mathsf{R}}})\,\mathsf{P} + \beta_0(1{+}\nu)\Gamma = -\omega_\rho\,\mathsf{P} + \beta_\rho\Sigma + \Xi_\rho, \quad [15]$$

$$\partial_{\mathsf{T}}\overset{\leftrightarrow}{\mathsf{S}} + \frac{1}{2}(\vec{\mathsf{R}}\cdot\partial_{\vec{\mathsf{R}}})\overset{\leftrightarrow}{\mathsf{S}} = -\omega_s\overset{\leftrightarrow}{\mathsf{S}} + \beta_s\overset{\leftrightarrow}{\Sigma}_d + \overset{\leftrightarrow}{\Xi}_s. \quad [16]$$

Synthesis fluctuations are described by a white Gaussian noise, with isotropic $\Xi_\rho$ and anisotropic $\overset{\leftrightarrow}{\Xi}_s$ components, with decaying spatial correlations. It is such that $\langle\Xi_\rho\rangle = 0$ and $\langle\overset{\leftrightarrow}{\Xi}_s\rangle = \overset{\leftrightarrow}{0}$, $\langle\overset{\leftrightarrow}{\Xi}_s\Xi_\rho\rangle = \overset{\leftrightarrow}{0}$, and,

$$\langle\Xi(\mathsf{T}_1,\vec{\mathsf{R}}_1)\Xi(\mathsf{T}_2,\vec{\mathsf{R}}_2)\rangle = \mathsf{K}_{\rho\rho}\delta(\mathsf{T}_1-\mathsf{T}_2)g(|\vec{\mathsf{R}}|) \quad [17]$$

$$\langle\overset{\leftrightarrow}{\Xi}(\mathsf{T}_1,\vec{\mathsf{R}}_1)\overset{\leftrightarrow}{\Xi}(\mathsf{T}_2,\vec{\mathsf{R}}_2)\rangle = \mathbf{K_{ss}}\delta(\mathsf{T}_1-\mathsf{T}_2)g(|\vec{\mathsf{R}}|), \quad [18]$$

where the Cartesian coordinates of $\mathbf{K_{ss}}$ are $\mathbf{K_{ss}}_{abcd} = \mathsf{K}_{ss}\left\{\delta_{ad}\delta_{bc}+\delta_{ac}\delta_{bd}-\delta_{ab}\delta_{cd}\right\}$ ($\delta$ is the Kronecker delta) and $g(x)$ decays quickly to 0 as $x>1$. (In practice, we use $g(x)=e^{-x}$.)

Equations [11-16], together with the form of the noise [17-18], amount to a system of 3 partial differential equations for velocity, $\vec{V}$, density, $\mathsf{P}$, and nematic tensor, $\overset{\leftrightarrow}{S}$, which should be supplemented with the boundary conditions $\overset{\leftrightarrow}{\Sigma}\cdot\vec{N}=\vec{0}$ where $N$ is the unit vector normal to the boundary.

#### RESPONSE TO PERTURBATIONS IN SYNTHESIS

##### General results

We here determine the explicit dependence of strain rate on fluctuations in synthesis. We consider the tissue size to be much larger than the correlation length, $\ell$, and than the characteristic scale of the perturbation. We assume that the perturbations do not induce large scale rotation ($\langle\partial_{\vec{\mathsf{R}}}\wedge\vec{\mathsf{V}}\rangle=\vec{0}$) and we chose the reference frame with origin such that $\vec{\mathsf{V}}(\mathsf{T},\vec{\mathsf{R}}=\vec{0})=\vec{0}$. Under these conditions and in the limit of infinite system size, the boundary conditions for stress are satisfied and Green's functions for the density, nematic tensor and flow velocity asymptotically vanish at infinity.

Noting that advective derivatives in Eqs. [15-16] expand spatial patterns at a rate 1/2, we introduce a modified Fourier transform such that the wavenumber is contracted by a factor $\exp(\mathsf{T}/2)$,

$$\widetilde{\Phi}(\mathsf{T},\vec{q}) = \int d^2\vec{\mathsf{R}}\exp(-\mathsf{T})\exp\left[-i(\vec{q}\cdot\vec{\mathsf{R}})\exp(-\mathsf{T}/2)\right]\Phi(\mathsf{T},\vec{\mathsf{R}}). \quad [19]$$

With this definition, the time-derivative of the Fourier transform $\widetilde{\Phi}$ is the Fourier transform of $\left[\partial_{\mathsf{T}}+\frac{1}{2}\vec{\mathsf{R}}\cdot\partial_{\vec{\mathsf{R}}}\right]\Phi(\mathsf{T},\vec{\mathsf{R}})$. Accordingly, the Fourier transforms of [15] and [16] are

$$\partial_{\mathsf{T}}\widetilde{\mathsf{P}} + \beta_0(1+\nu)\widetilde{\Gamma} = -\omega_\rho\widetilde{\mathsf{P}} + \beta_\rho\widetilde{\Sigma} + \widetilde{\Xi}_\rho, \quad [20]$$

$$\partial_{\mathsf{T}}\widetilde{\overset{\leftrightarrow}{\mathsf{S}}} = -\omega_s\widetilde{\overset{\leftrightarrow}{\mathsf{S}}} + \beta_s\widetilde{\overset{\leftrightarrow}{\Sigma}}_d + \widetilde{\overset{\leftrightarrow}{\Xi}}_s. \quad [21]$$

The Fourier transform of the momentum equation [11] is

$$\left[(1+\nu)\widetilde{\Gamma}+\widetilde{\mathsf{P}}\right]i\vec{q} + i\vec{q}\cdot\left[(1-\nu)\widetilde{\overset{\leftrightarrow}{\Gamma}}_d + \widetilde{\overset{\leftrightarrow}{\mathsf{S}}}\right] = \vec{0}. \quad [22]$$

Isotropic and deviatoric strain rate, which have 1 and 2 independent components, respectively, are not independent because they derive from the flow velocity (2 components) [12]: $\widetilde{\Gamma}=\widetilde{\Gamma}_{d\,qq}$, where we use the notation $\tilde{\Phi}_{qq}=\hat{q}\cdot\widetilde{\overset{\leftrightarrow}{\Phi}}\cdot\hat{q}$ for the $(\hat{q},\hat{q})$ component of the Fourier transform of tensor $\Phi$, $\hat{q}$ being the unit vector in the direction of the vector $q$. Using the consistency relation for strain rate, Eq. [22] is decomposed into isotropic and deviatoric components:

$$\widetilde{\Gamma}(\mathsf{T},\vec{q}) = -\frac{1}{2}[\widetilde{\mathsf{P}}(\mathsf{T},\vec{q})+\widetilde{\mathsf{S}}_{qq}(\mathsf{T},\vec{q})], \quad [23]$$

$$\overleftrightarrow{\widetilde{\mathsf{\Gamma}}}_d(\mathsf{T},\vec{q}) = \widetilde{\mathsf{\Gamma}}(\mathsf{T},\vec{q})[2\hat{q}\hat{q} - \overleftrightarrow{1}] - \frac{1}{1-\nu}[\hat{q}\vec{\widetilde{\mathsf{S}}}_{q\perp}(\mathsf{T},\vec{q}) + \vec{\widetilde{\mathsf{S}}}_{q\perp}(\mathsf{T},\vec{q})\hat{q}]. \qquad [24]$$

We then rewrite Eqs. [20-21] by combining the constitutive equations [13-14] and Eqs. [23-24] above, which yields ordinary differential equations for the Fourier transforms of density and nematic tensor,

$$\partial_{\mathsf{T}}\,\widetilde{\mathsf{P}} = -\left(\omega_\rho - \beta_0 - \frac{1-\nu}{2}(\beta_\rho - \beta_0)\right)\widetilde{\mathsf{P}} - \frac{1+\nu}{2}(\beta_\rho - \beta_0)\widetilde{\mathsf{S}}_{qq} + \widetilde{\Xi}_\rho, \qquad [25]$$

$$\partial_{\mathsf{T}}\,\overleftrightarrow{\widetilde{\mathsf{S}}} = -\omega_s\overleftrightarrow{\widetilde{\mathsf{S}}} + \beta_s\frac{1+\nu}{2}\widetilde{\mathsf{S}}_{qq}[2\hat{q}\hat{q} - \overleftrightarrow{1}] - \beta_s\frac{1-\nu}{2}[2\hat{q}\hat{q} - \overleftrightarrow{1}] + \overleftrightarrow{\widetilde{\Xi}}_s. \qquad [26]$$

The tensor $\overleftrightarrow{\widetilde{\mathsf{S}}}$ can then be written in terms of $\widetilde{\mathsf{S}}_{qq}$ and $\vec{\widetilde{\mathsf{S}}}_{q\perp}$, using the notation $\tilde{\Phi}_{qq} = \hat{q}\cdot\overleftrightarrow{\Phi}\cdot\hat{q}_\perp$, where $\hat{q}_\perp$ is the unit vector perpendicular to $\hat{q}$): $\overleftrightarrow{\widetilde{\mathsf{S}}} = \widetilde{\mathsf{S}}_{qq}[2\hat{q}\hat{q} - \overleftrightarrow{1}] + \{\hat{q}\vec{\widetilde{\mathsf{S}}}_{q\perp} + \vec{\widetilde{\mathsf{S}}}_{q\perp}\hat{q}\}$. It is therefore sufficient to know $\widetilde{\mathsf{S}}_{qq}$ and $\vec{\widetilde{\mathsf{S}}}_{q\perp}$ to fully determine $\overleftrightarrow{\widetilde{\mathsf{S}}}$.

The dynamics of $\widetilde{\mathsf{P}}$ and $\overleftrightarrow{\widetilde{\mathsf{S}}}$ is then described by

$$\partial_{\mathsf{T}}\begin{bmatrix}\widetilde{\mathsf{P}}(\mathsf{T},\vec{q})\\ \widetilde{\mathsf{S}}_{qq}(\mathsf{T},\vec{q})\end{bmatrix} = -\left[\omega\right]\cdot\begin{bmatrix}\widetilde{\mathsf{P}}(\mathsf{T},\vec{q})\\ \widetilde{\mathsf{S}}_{qq}(\mathsf{T},\vec{q})\end{bmatrix}, \qquad [27]$$

$$\partial_{\mathsf{T}}\vec{\widetilde{\mathsf{S}}}_{q\perp}(\mathsf{T},\vec{q}) = -\omega_s\vec{\widetilde{\mathsf{S}}}_{q\perp}(\mathsf{T},\vec{q}), \qquad [28]$$

where the relaxation matrix $\left[\omega\right]$ is given by

$$[\omega] = \begin{bmatrix}\omega_\rho - \beta_0 - \frac{1-\nu}{2}(\beta_\rho - \beta_0) & \frac{1+\nu}{2}(\beta_\rho - \beta_0)\\ \frac{1-\nu}{2}\beta_s & \omega_s - \frac{1+\nu}{2}\beta_s\end{bmatrix} \qquad [29]$$

It can be re expressed in terms of its eigenvalues $\omega_-$ and $\omega_-$ as $[\omega] = [P_+]\,\omega_+ + [P_-]\,\omega_-$ with $[P_\pm] = \begin{bmatrix}C_\rho^\pm \mp A & \pm A'\\ \pm A & C_s^\pm \mp A'\end{bmatrix}$ and $\omega_\pm$, $C_\varphi^\pm$, $A$, and $A'$ defined in Table I,

| Constant | Formula |
|---|---|
| $\omega_T$ | $\omega_s + \omega_\rho - \beta_0$ |
| $\Delta\omega$ | $\omega_\rho - \beta_0 - \omega_s$ |
| $B$ | $\frac{1-\nu}{2}(\beta_\rho - \beta_0) + \frac{1+\nu}{2}\beta_s$ |
| $\Delta\beta$ | $\frac{1-\nu}{2}(\beta_\rho - \beta_0) - \frac{1+\nu}{2}\beta_s$ |
| $\omega_\pm$ | $\frac{\omega_T - B}{2} \pm \frac{1}{2}\sqrt{\Delta\omega^2 + B^2 - 2\Delta\omega\Delta\beta}$ |
| $C_\rho^\pm$ | $\frac{1}{2} \pm \frac{1}{2}\frac{\Delta\omega + B - 2\Delta\beta}{\sqrt{\Delta\omega^2 + B^2 - 2\Delta\omega\Delta\beta}}$ |
| $C_s^\pm$ | $\frac{1}{2} \pm \frac{1}{2}\frac{-\Delta\omega + B + 2\Delta\beta}{\sqrt{\Delta\omega^2 + B^2 - 2\Delta\omega\Delta\beta}}$ |
| $A$ | $\frac{1}{2}\frac{B - \Delta\beta}{\sqrt{\Delta\omega^2 + B^2 - 2\Delta\omega\Delta\beta}}$ |
| $A'$ | $\frac{1}{2}\frac{B + \Delta\beta}{\sqrt{\Delta\omega^2 + B^2 - 2\Delta\omega\Delta\beta}}$ |

TABLE I. Values of the main constants appearing in calculations.

The solutions of [27] and [28] are

$$\begin{bmatrix}\widetilde{\mathsf{P}}(\mathsf{T},\vec{q})\\ \widetilde{\mathsf{S}}_{qq}(\mathsf{T},\vec{q})\end{bmatrix} = \int_{-\infty}^{\mathsf{T}} d\tau\left[e^{(\tau-\mathsf{T})\,[\omega]}\right]\cdot\begin{bmatrix}\widetilde{\Xi}_\rho(\tau,\vec{q})\\ \widetilde{\Xi}_{s\,qq}(\tau,\vec{q})\end{bmatrix}, \qquad [30]$$

$$\vec{\widetilde{\mathsf{S}}}_{q\perp}(\mathsf{T},\vec{q}) = \int_{-\infty}^{\mathsf{T}} d\tau\, e^{\omega_s(\tau-\mathsf{T})}\vec{\widetilde{\Xi}}_{s\,q\perp}(\tau,\vec{q}). \qquad [31]$$

Here $\left[e^{\tau\,[\omega]}\right]$ is the matrix exponential of $[\omega]$, which can be written as $\left[e^{\tau\,[\omega]}\right] = [P_+]\, e^{\omega_+\tau} + [P_-]\, e^{\omega_-\tau}$.

Finally, the strain rate, $\overset{\leftrightarrow}{\widetilde{\Gamma}}$, has two independent components, $\Gamma$ and $\vec{\Gamma}_{q\perp}$; accordingly $\overset{\leftrightarrow}{\widetilde{\Gamma}} = \widetilde{\Gamma}\hat{q}\hat{q} + \hat{q}\vec{\widetilde{\Gamma}}_{q\perp} + \vec{\widetilde{\Gamma}}_{q\perp}\hat{q}$. The two independent components can be derived from [23] and [24] by substituting density and nematic tensor in in [30] and [31]:

$$\widetilde{\Gamma}(\mathsf{T},\vec{q}) = -\int_{-\infty}^{\mathsf{T}} d\tau \sum_{\varepsilon\in\{-,+\}} \frac{e^{\omega_\varepsilon(\tau-\mathsf{T})}}{2} \left[C^{\varepsilon}_{\rho}\, \widetilde{\Xi}_{\rho}(\tau,\vec{q}) + C^{\varepsilon}_{s}\, \widetilde{\Xi}_{s\,qq}(\tau,\vec{q})\right], \tag{32}$$

$$\vec{\widetilde{\Gamma}}_{q\perp}(\mathsf{T},\vec{q}) = -\frac{1}{1-\nu}\int_{-\infty}^{\mathsf{T}} d\tau e^{\omega_s(\tau-\mathsf{T})}\, \vec{\widetilde{\Xi}}_{s\,q\perp}(\tau,\vec{q}). \tag{33}$$

$\omega_\pm$ and $C^{\pm}_{\varphi}$ are defined in the Table I. These two equations yield the fluctuations in strain rate whenever the fluctuations in synthesis are given. Throughout this work, we assume that uniform state is stable, which holds for $\omega_s > 0$ and $\omega_\pm > 0$.

##### Example: disk-shaped isotropic perturbation

We consider a perturbation of isotropic synthesis localised at time $t = 0$ and inside a disk of radius $R = 1$, $\Xi_\rho(\mathsf{T},\vec{\mathsf{R}}) = \delta(\mathsf{T})\mathrm{H}(1-|\vec{\mathsf{R}}|)$, where $H$ is the Heaviside function), and $\overset{\leftrightarrow}{\Xi}_s(\mathsf{T},\vec{\mathsf{R}}) = \overset{\leftrightarrow}{0}$. We first substitute the Fourier transform of $\Xi_\rho$ and $\overset{\leftrightarrow}{\Xi}_s$ in [30], [31], [32] and [33]. The tension is then deduced from [13] and [14]. Finally, we obtain all fields by computing the inverse Fourier transform $\Phi(\mathsf{T},\vec{\mathsf{R}}) = \int \frac{d^2\vec{q}e^{\mathsf{T}}}{(2\pi)^2} e^{i(\vec{q}\cdot\vec{\mathsf{R}})e^{\frac{\mathsf{T}}{2}}}\widetilde{\Phi}(\mathsf{T},\vec{q})$. All fields take the generic form,

$$\Phi(\mathsf{T},\vec{\mathsf{R}}) = \mathcal{A}_\Phi(\mathsf{T})\,\mathcal{B}_\Phi(\vec{\mathsf{R}}\,e^{-\frac{\mathsf{T}}{2}}), \quad \Phi\in\{\mathsf{P},\,\overset{\leftrightarrow}{\mathsf{S}},\,\overset{\leftrightarrow}{\Sigma},\,\overset{\leftrightarrow}{\Gamma}\},$$

where $\mathcal{A}_\Phi$ and $\mathcal{B}_\Phi$ are, respectively, the amplitude and the spatial rescaled form of the field $\Phi$. We find:

$$\mathcal{A}_{\mathsf{P}}(\mathsf{T}) = (C^{+}_{\rho} - A)\,e^{-\omega_+\mathsf{T}} + (C^{-}_{\rho} + A)\,e^{-\omega_-\mathsf{T}}, \tag{34}$$

$$\mathcal{A}_{\mathsf{S}}(\mathsf{T}) = -A\,e^{-\omega_+\mathsf{T}} + A\,e^{-\omega_-\mathsf{T}}, \tag{35}$$

$$\mathcal{A}_{\Gamma}(\mathsf{T}) = -\frac{1}{2}\left\{C^{+}_{\rho}\,e^{-\omega_+\mathsf{T}} + C^{-}_{\rho}\,e^{-\omega_-\mathsf{T}}\right\}, \tag{36}$$

$$\mathcal{A}_{\Sigma}(\mathsf{T}) = \left(\frac{1-\nu}{2}C^{+}_{\rho} - A\right)\,e^{-\omega_+\mathsf{T}} + \left(\frac{1-\nu}{2}C^{-}_{\rho} + A\right)\,e^{-\omega_-\mathsf{T}}; \tag{37}$$

$$\mathcal{B}_{\mathsf{P}}(\vec{\mathsf{R}}) = \mathrm{H}(1-|\vec{\mathsf{R}}|), \tag{38}$$

$$\mathcal{B}_{\mathsf{S}}(\vec{\mathsf{R}}) = [2\,\hat{R}\hat{R} - \overset{\leftrightarrow}{1}]\,\mathrm{F}(|\vec{\mathsf{R}}|), \tag{39}$$

$$\mathcal{B}_{\Gamma}(\vec{\mathsf{R}}) = \left[\overset{\leftrightarrow}{1}\,\mathrm{H}(1-|\vec{\mathsf{R}}|) - [2\,\hat{R}\hat{R} - \overset{\leftrightarrow}{1}]\,\mathrm{F}(|\vec{\mathsf{R}}|)\right], \tag{40}$$

$$\mathcal{B}_{\Sigma}(\vec{\mathsf{R}}) = \left[\overset{\leftrightarrow}{1}\,\mathrm{H}(1-|\vec{\mathsf{R}}|) + [2\,\hat{R}\hat{R} - \overset{\leftrightarrow}{1}]\,\mathrm{F}(|\vec{\mathsf{R}}|)\right], \tag{41}$$

where $\hat{R} = \vec{\mathsf{R}}/|\vec{\mathsf{R}}|$, and $F$ is defined by $F(x) = 0$ if $x \le 1$ and $F(x) = 1/x^2$ if $x > 1$.

We find that all fields decay to 0. In most cases(? ), this occurs without oscillations (over-damped relaxation) when $\beta_s < \beta_s^{\mathrm{t}}$ and with oscillations (under-damped relaxation) when $\beta_s > \beta_s^{\mathrm{t}}$ (we named the corresponding regimes hypo- and hyper-responsive, respectively), the threshold value of $\beta_s$ being

$$\beta_s^{\mathrm{t}} = \frac{1}{1+\nu}\left\{-2\Delta\omega - (1-\nu)(\beta_\rho - \beta_0) - 2\sqrt{2\Delta\omega(1-\nu)(\beta_\rho - \beta_0)}\right\}, \qquad [42]$$

where $\Delta\omega$ is defined in Table I.

The characteristic decay time $\tau_c$ is the maximal relaxation time scale in response to a perturbation. It reads $\tau_c = 1/\mathrm{Re}(\omega_-)$ if deviatoric noise in synthesis is absent and $\tau_c = \max\{1/\mathrm{Re}(\omega_-), 1/\omega_s\}$ otherwise (the constants are defined in Table I). The first case is due to some components of the nematic tensor relaxing independently of the density. In either case, $\tau_c$ depends on the kinetics of synthesis and mechanical feedbacks.

All these results are discussed in the main text of the article, with graphs represented for the parameters $\{\omega_\rho, \omega_s, \beta_0, \nu\}$ equal to $\{1, 1, 1, 0\}$.

#### GROWTH FLUCTUATIONS

##### Correlation functions

Using [32] and [33], the rescaled velocity $\vec{\mathsf{V}}(\mathsf{T},\vec{\mathsf{R}})$ can be written as a function of $\Xi_\rho(\mathsf{T},\vec{\mathsf{R}})$ and $\overset{\leftrightarrow}{\Xi}_s(\mathsf{T},\vec{\mathsf{R}})$, taking the form

$$\begin{aligned}\vec{\mathsf{V}}(\mathsf{T},\vec{\mathsf{R}}) = &-\frac{1}{2\pi}\int_{-\infty}^{\mathsf{T}} d\tau\, e^{(-\frac{1}{2}+\omega_\varepsilon)(\tau-\mathsf{T})}\int\frac{d^2\vec{u}}{|\vec{u}|}\left\{\hat{u}\, C_\rho^\varepsilon\left[\Xi_\rho(\tau,\vec{u}) - \Xi_\rho(\tau,\vec{u}+\vec{\mathsf{R}}\,e^{\frac{\mathsf{T}}{2}})\right] + C_s^\varepsilon\left[\vec{\Xi}_{s\,u\perp}(\tau,\vec{u}) - \vec{\Xi}_{s\,u\perp}(\tau,\vec{u}+\vec{\mathsf{R}}\,e^{\frac{\mathsf{T}}{2}})\right]\right\}\\ &-\frac{1}{\pi(1-\nu)}\int_{-\infty}^{\mathsf{T}} d\tau\, e^{(-\frac{1}{2}+\omega_s)(\tau-\mathsf{T})}\int\frac{d^2\vec{u}}{|\vec{u}|}\,\hat{u}\left[\Xi_{s\,uu}(\tau,\vec{u}) - \Xi_{s\,uu}(\tau,\vec{u}+\vec{\mathsf{R}}\,e^{\frac{\mathsf{T}}{2}})\right], \qquad [43]\end{aligned}$$

where we used the assumption of no global rotation and no translation of the origin of the coordinate system. We derive the correlation function $\langle\vec{\mathsf{V}}(\mathsf{T},\vec{\mathsf{R}}_1)\vec{\mathsf{V}}(\mathsf{T},\vec{\mathsf{R}}_2+\Delta\mathsf{T})\rangle$ by combining [43] with [17-18]. This derivation, conceptually simple but tedious, involves the calculation of various integrals, which are summarised below.

$$\begin{aligned}&\int\frac{d^2\vec{u}}{|\vec{u}|}\int\frac{d^2\vec{w}}{|\vec{w}|}\left\langle\left[\vec{\Phi}(0,\vec{u}) - \vec{\Phi}(0,\vec{u}+\vec{\mathsf{R}})\right]\left[\vec{\Phi}(\tau,\vec{w}) - \vec{\Phi}(\tau,\vec{w}+\vec{y})\right]\right\rangle \qquad [44]\\ &\qquad = 2\pi^2\overset{\leftrightarrow}{1}\delta(\mathsf{T})\left\{\int_0^{|\vec{\mathsf{R}}|}\frac{dR}{R}\int_0^R dr\,r\,g(r) + \int_0^{|\vec{y}|}\frac{dR}{R}\int_0^R dr\,r\,g(r) - \int_0^{|\vec{\mathsf{R}}-\vec{y}|}\frac{dR}{R}\int_0^R dr\,r\,g(r)\right\}\end{aligned}$$

where the vector field $\vec{\Phi}(\tau,\vec{u})$ can be $\hat{u}\,\Xi_\rho(\tau,\vec{u})$, $\vec{\Xi}_{s\,u\perp}(\tau,\vec{u})$, or $\hat{u}\,\Xi_{s\,uu}(\tau,\vec{u})$, which is a consequence of the relation

$$2\pi^2\overset{\leftrightarrow}{1}\int_0^{|\vec{\mathsf{R}}|}\frac{dR}{R}\int_0^R dr\,r\,g(r) = \int\frac{d^2\vec{u}}{|\vec{u}|}\int\frac{d^2\vec{w}}{|\vec{w}|}\left\langle\vec{\Phi}(\tau,\vec{u})\vec{\Phi}(\tau,\vec{w})\right\rangle - \left\langle\vec{\Phi}(\tau,\vec{u})\vec{\Phi}(\tau,\vec{w}+\vec{\mathsf{R}})\right\rangle.$$

Other integrals vanish, such as for

$$\int\frac{d^2\vec{u}}{|\vec{u}|}\int\frac{d^2\vec{w}}{|\vec{w}|}\left\langle\left[\vec{\Phi}(\tau,\vec{u}) - \vec{\Phi}(\tau,\vec{u}+\vec{\mathsf{R}})\right]\left[\vec{\Psi}(\tau,\vec{w}) - \vec{\Psi}(\tau,\vec{w}+\vec{y})\right]\right\rangle = \overset{\leftrightarrow}{0},$$

where the two different vector fields $\vec{\Phi}(\tau,\vec{u})$ and $\vec{\Psi}(\tau,\vec{u})$ belong to $\{\hat{u}\,\Xi_\rho(\tau,\vec{u}), \vec{\Xi}_{s\,u\perp}(\tau,\vec{u}), \hat{u}\,\Xi_{s\,uu}(\tau,\vec{u})\}$, which is a consequence of $\langle\Xi_\rho\overset{\leftrightarrow}{\Xi}_s\rangle = \overset{\leftrightarrow}{0}$ and of

$$\overset{\leftrightarrow}{0} = \int\frac{d^2\vec{u}}{|\vec{u}|}\hat{u}\int\frac{d^2\vec{w}}{|\vec{w}|}\left\langle\Xi_{s\,uu}(\tau,\vec{u})\vec{\Xi}_{s\,u\perp}(\tau,\vec{w})\right\rangle - \left\langle\Xi_{s\,uu}(\tau,\vec{u})\vec{\Xi}_{s\,u\perp}(\tau,\vec{w}+\vec{y})\right\rangle.$$

The correlation function for the velocity derives from [43-44] and the integrals above:

$$\langle\vec{\mathsf{V}}(\mathsf{T},\vec{\mathsf{R}}_1)\vec{\mathsf{V}}(\mathsf{T},\vec{\mathsf{R}}_2+\Delta\mathsf{T})\rangle = \overset{\leftrightarrow}{1}\left\{\mathsf{K}_{\varphi_1\varphi_2}C_{\varphi_1}^{\varepsilon_1}C_{\varphi_2}^{\varepsilon_2}\mathcal{I}(\vec{\mathsf{R}}_1,\vec{\mathsf{R}}_2,\Delta\mathsf{T},\omega_{\varepsilon_1},\omega_{\varepsilon_2}) + \frac{4\mathsf{K}_{ss}}{(1-\nu)^2}\mathcal{I}(\vec{\mathsf{R}}_1,\vec{\mathsf{R}}_2,\Delta\mathsf{T},\omega_s,\omega_s)\right\}, \qquad [45]$$

with $\Delta\mathsf{T} > 0$ and,

$$\mathcal{I}(\vec{\mathsf{R}}_1, \vec{\mathsf{R}}_2, \Delta\mathsf{T}, \omega_1, \omega_2) = e^{\Delta\mathsf{T}(\frac{1}{2}-\omega_1)} \left\{ I(|\vec{\mathsf{R}}_2| e^{-\frac{\Delta\mathsf{T}}{2}}, \omega_1+\omega_2) + I(|\vec{\mathsf{R}}_1|, \omega_1+\omega_2) - I(|\vec{\mathsf{R}}_1 - \vec{\mathsf{R}}_2 e^{-\frac{\Delta\mathsf{T}}{2}}|, \omega_1+\omega_2) \right\},$$

where $I(x,\omega) = \frac{1}{2}\int_{-\infty}^{0} d\tau e^{\tau(\omega-1)} \int_0^{x\, e^{\frac{\tau}{2}}} \frac{dR}{R} \int_0^R dr r\, g(r)$ can be rewritten via multiple integrations by parts as

$$I(x,\omega_T) = \frac{1}{2}\left\{ \frac{\log(x) - \frac{1}{2\omega_T-2}}{\omega_T - 1}[1-(1+x)e^{-x}] - \frac{\int_0^x du u \log(u) e^{-u}}{\omega_T-1} + x^{2-2\omega_T} \frac{\int_0^x du u^{2\omega_T-1} e^{-u}}{2(\omega_T-1)^2} \right\}, \quad [46]$$

assuming $g(x) = e^{-x}$.

From this, we may derive the correlation function of the areal strain rate:

$$\langle \Gamma(\mathsf{T}, \vec{\mathsf{R}}_1)\Gamma(\mathsf{T}+\Delta\mathsf{T}, \vec{\mathsf{R}}_2)\rangle = \mathsf{K}_{\varphi_1\varphi_2} C_{\varphi_1}^{\varepsilon_1} C_{\varphi_2}^{\varepsilon_2} e^{-\Delta\mathsf{T}\omega_{\varepsilon_1}} I_2(|\vec{\mathsf{R}}_1 - \vec{\mathsf{R}}_2 e^{-\frac{\Delta\mathsf{T}}{2}}|, \omega_{\varepsilon_1}+\omega_{\varepsilon_2}) + \frac{4\mathsf{K}_{ss}}{(1-\nu)^2} e^{-\Delta\mathsf{T}\omega_s} I_2(|\vec{\mathsf{R}}_1 - \vec{\mathsf{R}}_2 e^{-\frac{\Delta\mathsf{T}}{2}}|, 2\omega_s), \quad [47]$$

with

$$I_2(x,\omega) = \frac{1}{4x^{2\omega}} \int_0^x du u^{2\omega-1} g(u)$$

. The correlation function of the areal strain rate has power-law tails, $\langle \Gamma(\vec{\mathsf{R}}_1)\Gamma(\vec{\mathsf{R}}_2)\rangle \sim \|\vec{\mathsf{R}}_1 - \vec{\mathsf{R}}_2\|^{-4/\tau_c}$, yielding long-range correlations.

#### Results

In the main text of the article, we defined time-correlations of growth at scale $\mathsf{R}$, $G(\mathsf{T},\mathsf{R})$, as the time-correlation of growth in a disk of radius $\mathsf{R}$. This coarse-grained time correlation is related to the velocity correlation function through

$$G(\mathsf{T},\mathsf{R}) = \frac{1}{(\pi\,\mathsf{R}^2)^2} \left\langle \int_0^{2\pi} d\theta \int_0^{2\pi} d\varphi \left(\vec{\mathsf{R}}(\theta)\cdot\vec{\mathsf{V}}(\vec{\mathsf{R}}(\theta),0)\right)\left(\vec{\mathsf{R}}(\varphi)\cdot\vec{\mathsf{V}}(\vec{\mathsf{R}}(\varphi),\mathsf{T})\right)\right\rangle.$$

We use [45] to compute $G$, assuming $\mathsf{R} \gg 1$,

$$G(\mathsf{T},\mathsf{R}) \overset{\mathsf{R}\gg 1}{=} \mathsf{K}_{\varphi_1\varphi_2} C_{\varphi_1}^{\varepsilon_1} C_{\varphi_2}^{\varepsilon_2}\, e^{-\omega_{\varepsilon_1}\mathsf{T}} \mathcal{S}(\mathsf{T},\mathsf{R},\omega_{\varepsilon_1}+\omega_{\varepsilon_2}) + \frac{4\mathsf{K}_{ss}}{(1-\nu)^2}\, e^{-\omega_s\mathsf{T}} \mathcal{S}(\mathsf{T},\mathsf{R},2\omega_s) \quad [48]$$

with,

$$\mathcal{S}(\mathsf{T},\mathsf{R},\omega_T) = \left\{ \frac{1}{\mathsf{R}^2}\frac{1}{\omega_T-1} + \frac{s(\mathsf{T},\omega_T)}{\mathsf{R}^{2\omega_T}} \frac{\Gamma_e(2\omega_T)}{1-\omega_T} \right\}$$

where

$$s(\mathsf{T},\omega) = \frac{e^{\frac{\mathsf{T}}{2}}}{2\pi(1-\omega)} \int_0^{2\pi} d\theta (1+e^{-\mathsf{T}} + 2e^{-\frac{\mathsf{T}}{2}}\cos\theta)^{1-\omega}.$$

These results are discussed in in the main text of the article. In [48], $G$ depends on $\mathsf{R}$ only through a linear combination of power laws $\mathsf{R}^\alpha$ where $\alpha \in \{-2, -4\omega_+, -4\omega_-, -2\omega_+ - 2\omega_-, -4\omega_s\}$, $\omega_\pm$ and $\omega_s$ being the eigenvalues of the relaxation matrix. Hence $\alpha \in \{-2, -4/\tau_c\}$ because the correlation time $\tau_c$ is the maximal relaxation time scale $\tau_c = \max\{1/\mathrm{Re}(\omega_-), 1/\omega_s\}$. We thus find that, when $\tau_c < 2$, growth mean square deviation scales with the inverse of the coarse-graining area, $G(\mathsf{R},0) \propto \mathsf{R}^{-2}$, an exponent that corresponds to the central limit theorem. When $\tau_c > 2$, growth mean square deviation decays more slowly with the coarse-graining area, $G(\mathsf{R},0) \propto \mathsf{R}^{-4/\tau_c}$.

This second scaling can be rationalized as follows. Growth fluctuations, $G$, of a surface of size $\mathsf{R}$ are not much influenced by recent synthesis fluctuations whenever $\mathsf{R} \gg 1$, because these fluctuations are averaged over many correlations lengths. But $G$ retains the memory of fluctuations in synthesis at an earlier time $\mathsf{T} - \Delta\mathsf{T}$ when the surface size, $\mathsf{R}\, e^{-\Delta\mathsf{T}/2}$, was comparable to the correlation length (1 in dimensionless units). $G$ responds to these earlier fluctuations with a relaxation time $\tau_c$, $G \sim e^{-2\frac{\Delta\mathsf{T}}{\tau_c}} \sim \mathsf{R}^{-\frac{4}{\tau_c}}$ by elimination of $\Delta\mathsf{T}$.

#### FLUCTUATIONS OF ORGAN SHAPE

To characterise fluctuations at organ scale, we consider the fluctuations of the relative position between two material points of the organ. Because of invariance by translation, it suffices to choose their initial positions to be the origin of coordinate system and a vector $\vec{X}_0$. We therefore consider the Lagrangian position of a material point $\vec{X}$ of the tissue initially at position $\vec{X}_0$,

$$\frac{d\vec{X}}{d\mathsf{T}} = \frac{\vec{X}(\mathsf{T})}{2} + \vec{\mathsf{V}}(\mathsf{T}, \vec{X}(\mathsf{T})), \qquad \vec{X}(0) = \vec{X}_0. \qquad [49]$$

We hereafter compute the statistical properties of the vector $\vec{X}$ from growth rate distributions.

##### Distribution

The probability for $\vec{X}(\mathsf{T})$ has the general expression

$$\mathcal{P}\left[\vec{X}(\mathsf{T})\right] = \left\langle \delta(\vec{X}(0) - \vec{X}_0)\, \delta\left[\frac{d\vec{X}(\mathsf{T})}{d\mathsf{T}} - \frac{\vec{X}(\mathsf{T})}{2} - \vec{\mathsf{V}}(\mathsf{T}, \vec{X}(\mathsf{T}))\right] \right\rangle. \qquad [50]$$

where $\delta\left[\cdot\right]$ is a delta functional that can be defined as

$$\delta\left[\cdot\right] \equiv \int \mathcal{D}\left[\vec{q}\right] e^{i \int\, d\mathsf{T}\, \vec{q}(\mathsf{T}) \cdot \{\cdot\}(\mathsf{T})},$$

with $\int \mathcal{D}[\vec{q}]$ being the functional integral over the space of functions of $\vec{q}$. $\vec{\mathsf{V}}$ has a Gaussian distribution with kernel

$$\mathcal{P}[\vec{\mathsf{V}}] \propto e^{-\frac{1}{2}\int_0^\infty d\mathsf{T}_1 \int d^2\vec{\mathsf{R}}_1 \int_0^\infty d\mathsf{T}_2 \int d^2\vec{\mathsf{R}}_2\, \vec{\mathsf{V}}(\mathsf{T}_1,\vec{\mathsf{R}}_1) \overleftrightarrow{\mathcal{H}}(\mathsf{T}_1,\mathsf{T}_2,\vec{\mathsf{R}}_1,\vec{\mathsf{R}}_2) \vec{\mathsf{V}}(\mathsf{T}_2,\vec{\mathsf{R}}_2)}, \qquad [51]$$

where the Green function of $\overleftrightarrow{\mathcal{H}}(\mathsf{T}_1, \mathsf{T}_2, \vec{\mathsf{R}}_1, \vec{\mathsf{R}}_2)$ is $\langle \vec{\mathsf{V}}(\mathsf{T}_1, \vec{\mathsf{R}}_1)\vec{\mathsf{V}}(\mathsf{T}_2, \vec{\mathsf{R}}_2)\rangle$. The probability $\mathcal{P}\left[\vec{X}(\mathsf{T})\right] = \langle \int \mathcal{D}\left[\vec{q}\right] e^{i \int\, d\mathsf{T}\, \vec{q}(\mathsf{T}) \cdot \vec{X}(\mathsf{T})} \rangle$ can be explicitly written in terms of the probability $\mathcal{P}[\vec{\mathsf{V}}(\mathsf{T}, \vec{\mathsf{R}})]$ in [51], which yields

$$\mathcal{P}\left[\vec{X}(\mathsf{T})\right] \propto \delta(\vec{X}(0) - \vec{X}_0) \int \mathcal{D}\left[\vec{\mathsf{V}}\right] \int \mathcal{D}\left[\vec{q}\right] e^{i\int_0^\infty d\mathsf{T}\, \vec{q}(\mathsf{T}) \cdot \left\{ \frac{d\vec{X}(\mathsf{T})}{d\mathsf{T}} - \frac{\vec{X}(\mathsf{T})}{2} - \vec{\mathsf{V}}(\mathsf{T},\vec{X}(\mathsf{T})) \right\}} e^{-\int\int d\mathsf{T}_1 d\mathsf{T}_2 d^2\vec{\mathsf{R}}_1 d^2\vec{\mathsf{R}}_2 \vec{\mathsf{V}}(\mathsf{T}_1,\vec{\mathsf{R}}_1) \mathcal{H}(|\mathsf{T}_1,\mathsf{T}_2,\vec{\mathsf{R}}_1,\vec{\mathsf{R}}_2) \vec{\mathsf{V}}(\mathsf{T}_2,\vec{\mathsf{R}}_2)}, \qquad [52]$$

We recognize a Gaussian integral over $\vec{\mathsf{V}}$ in the r.h.s. of [52], which can be written as

$$\mathcal{P}\left[\vec{X}(\mathsf{T})\right] = \delta(\vec{X}(0) - \vec{X}_0) \int \mathcal{D}\left[\vec{q}\right] e^{i \int_0^\infty\, d\mathsf{T}\, \vec{q}(\mathsf{T}) \cdot \left\{ \frac{d\vec{X}(\mathsf{T})}{d\mathsf{T}} - \frac{1}{2}\vec{X}(\mathsf{T}) \right\}} e^{-\frac{1}{2}\int_0^\infty d\mathsf{T}_1 \int_0^\infty d\mathsf{T}_2\, \vec{q}(\mathsf{T}_1) \langle \vec{\mathsf{V}}(\mathsf{T}_1,\vec{X}(\mathsf{T}_1)) \vec{\mathsf{V}}(\mathsf{T}_2,\vec{X}(\mathsf{T}_2)) \rangle \vec{q}(\mathsf{T}_2)} \qquad [53]$$

We then make the change of variable $\mathsf{T}_1 = \mathsf{T}$, $\mathsf{T}_2 = \mathsf{T} + \tau$ in the last integral. We then consider time scales that are much bigger than the relaxation time, $T \gg \tau_c$, for which we make the approximation

$$\int_0^\infty d\mathsf{T}_1 \int_0^\infty d\mathsf{T}_2\, \vec{q}(\mathsf{T}_1) \langle \vec{\mathsf{V}}(\mathsf{T}_1, \vec{X}(\mathsf{T}_1)) \vec{\mathsf{V}}(\mathsf{T}_2, \vec{X}(\mathsf{T}_2)) \rangle \vec{q}(\mathsf{T}_2) \simeq \int_0^\infty d\mathsf{T} \int_{-\infty}^\infty d\tau\, \vec{q}(\mathsf{T}) \cdot \langle \vec{\mathsf{V}}(\mathsf{T}, \vec{X}(\mathsf{T})) \vec{\mathsf{V}}(\mathsf{T}+\tau, \vec{X}(\mathsf{T}+\tau)) \rangle \cdot \vec{q}(\mathsf{T}+\tau).$$

In the limit $\Gamma \ll 1$, as long as $\Gamma\tau_c \lesssim 1$, we may also make the approximations $\vec{X}(\mathsf{T}+\tau) \simeq \vec{X}(\mathsf{T}) e^{\frac{\tau}{2}}$ and $\vec{q}(\mathsf{T}+\tau) \simeq \vec{q}(\mathsf{T}) e^{-\frac{\tau}{2}}$ in this last expression. Altogether, Equation [53] becomes

$$\mathcal{P}\left[\vec{X}(\mathsf{T})\right] = \delta(\vec{X}(0) - \vec{X}_0) \int \mathcal{D}\left[\vec{q}\right] e^{i \int\, d\mathsf{T}\, \vec{q}(\mathsf{T}) \left\{ \frac{d\vec{X}(\mathsf{T})}{d\mathsf{T}} - \frac{1}{2}\vec{X}(\mathsf{T}) \right\}} e^{-\frac{1}{4}\int d\mathsf{T} |\vec{q}|^2(\mathsf{T}) \int_{-\infty}^\infty d\tau e^{-\frac{\tau}{2}} \langle \vec{\mathsf{V}}(\mathsf{T},\vec{X}(\mathsf{T})) \cdot \vec{\mathsf{V}}(\mathsf{T}+\tau, \vec{X}(\mathsf{T}) e^{\frac{\tau}{2}}) \rangle}, \qquad [54]$$

where we recognise again the Fourier transform of a Gaussian functional. More explicitly

$$\mathcal{P}\left[\vec{X}(\mathsf{T})\right] \propto \delta(\vec{X}(0) - \vec{X}_0) e^{-\mathcal{A}\left[\vec{X}(\mathsf{T})\right]}, \qquad [55]$$

with

$$\mathcal{A}\left[\vec{X}(\mathsf{T})\right] = \int d\mathsf{T}\, \frac{\left| \frac{d\vec{X}(\mathsf{T})}{d\mathsf{T}} - \frac{1}{2}\vec{X}(\mathsf{T}) \right|^2}{2\int_0^\infty d\tau e^{-\frac{\tau}{2}} \langle \vec{\mathsf{V}}(\mathsf{T}, \vec{X}(\mathsf{T})) \cdot \vec{\mathsf{V}}(\mathsf{T}+\tau, \vec{X}(\mathsf{T}) e^{\frac{\tau}{2}}) \rangle}. \qquad [56]$$

We hereafter apply the saddle point method to compute the first cumulants of $\mathcal{P}\left[\vec{X}(\mathsf{T})\right]$.

##### Cumulant generating functional and average displacements

We first write the cumulant generating functional $\mathcal{W}$ as the functional that associates each vector function $\vec{h}(T)$ to the quantity

$$\mathcal{W}\left[\vec{h}(\mathsf{T})\right] \equiv \log\left\langle \exp\left(\int d\mathsf{T}\,\vec{h}(\mathsf{T})\cdot\vec{X}(\mathsf{T})\right)\right\rangle .$$

In the limit when $\Gamma\to 0$, the saddle point method yields

$$\mathcal{W}\left[\vec{h}(\mathsf{T})\right] \simeq -\mathcal{A}\left[\vec{X}_e(\mathsf{T})\right] + \int d\mathsf{T}\,\vec{h}(\mathsf{T})\cdot\vec{X}_e(\mathsf{T}), \qquad [57]$$

where $\vec{X}_e$ depends of $\vec{h}$ through

$$\vec{h}(\mathsf{T}) = \frac{\delta\mathcal{A}}{\delta\vec{X}(\mathsf{T})}\left[\vec{X}_e\right]. \qquad [58]$$

The average displacement derives from cumulant generating functional,

$$\langle\vec{X}(\mathsf{T})\rangle = \frac{\delta\mathcal{W}}{\delta\vec{h}(\mathsf{T})}\left[0\right].$$

From [57], we therefore deduce

$$\langle\vec{X}(\mathsf{T})\rangle = -\int d\mathsf{T}'\frac{\delta\mathcal{A}}{\delta\vec{X}(\mathsf{T}')}\left[\vec{X}_e\right]\frac{\delta\vec{X}_e(\mathsf{T}')}{\delta\vec{h}(\mathsf{T})} + \int d\mathsf{T}'\,\vec{h}(\mathsf{T}')\cdot\frac{\delta\vec{X}_e(\mathsf{T}')}{\delta\vec{h}(\mathsf{T})} + \vec{X}_e(\mathsf{T}),$$

where the first two terms cancel out because of (58), leading to the average displacement vector

$$\langle\vec{X}(\mathsf{T})\rangle = \left.\vec{X}_e(\mathsf{T})\right|_{\left[\vec{h}=\vec{0}\right]} = \vec{X}_0\,e^{\frac{1}{2}\mathsf{T}}. \qquad [59]$$

This first result could easily be expected from the average strain rate.

##### Standard deviations

To characterize fluctuations, we compute the second moment of $\mathcal{P}\left[\vec{X}(\mathsf{T})\right]$, which is the second derivative of the cumulant generating functional:

$$\langle\Delta\vec{X}(\mathsf{T}_1)\Delta\vec{X}(\mathsf{T}_2)\rangle = \frac{\delta^2\mathcal{W}}{\delta\vec{h}(\mathsf{T}_1)\delta\vec{h}(\mathsf{T}_2)}\left[0\right] = \left.\frac{\delta\vec{X}_e(\mathsf{T}_1)}{\delta\vec{h}(\mathsf{T}_2)}\right|_{[h=0]},$$

where $\left.\frac{\delta\vec{X}_e(\mathsf{T}_1)}{\delta\vec{h}(\mathsf{T}_2)}\right|_{\left[\vec{h}=\vec{0}\right]}$ can be found by taking the functional derivative of [58] with respect to $\vec{h}$:

$$\int d\mathsf{T}'\,\frac{\delta\mathcal{A}}{\delta\vec{X}(\mathsf{T}_1)\delta\vec{X}(\mathsf{T}')}\left[\vec{X}_e\right]\cdot\left.\frac{\delta\vec{X}_e(\mathsf{T}')}{\delta\vec{h}(\mathsf{T}_2)}\right|_{\left[\vec{h}=\vec{0}\right]} = \delta(\mathsf{T}_1-\mathsf{T}_2) \qquad [60]$$

$\left.\frac{\delta\vec{X}_e(\mathsf{T}_1)}{\delta\vec{h}(\mathsf{T}_2)}\right|_{\left[\vec{h}=\vec{0}\right]}$ is therefore the Green's function of $\frac{\delta\mathcal{A}}{\delta\vec{X}(\mathsf{T}_1)\delta\vec{X}(\mathsf{T}_2)}\left[\vec{h}=\vec{0}\right]$. Starting from [56], we derive

$$\frac{\delta\mathcal{A}}{\delta\vec{X}(\mathsf{T}_1)\delta\vec{X}(\mathsf{T}_2)}\left[\vec{h}=\vec{0}\right] = \frac{\left[-\frac{d}{d\mathsf{T}_1}-\frac{1}{2}+\frac{dX_e}{d\mathsf{T}_1}\frac{\partial_{X_e}\int_0^\infty d\tau e^{-\frac{\tau}{2}}\langle\vec{\mathsf{V}}(\mathsf{T},\vec{X}_e(\mathsf{T}))\cdot\vec{\mathsf{V}}(\mathsf{T}+\tau,\vec{X}_e(\mathsf{T})e^{\frac{\tau}{2}})\rangle}{\int_0^\infty d\tau e^{-\frac{\tau}{2}}\langle\vec{\mathsf{V}}(\mathsf{T},\vec{X}_e(\mathsf{T}))\cdot\vec{\mathsf{V}}(\mathsf{T}+\tau,\vec{X}_e(\mathsf{T})e^{\frac{\tau}{2}})\rangle\rangle}\right]\left[\frac{d}{d\mathsf{T}_1}-\frac{1}{2}\right]}{\int_0^\infty d\tau e^{-\frac{\tau}{2}}\langle\vec{\mathsf{V}}(\mathsf{T},\vec{X}_e(\mathsf{T}))\cdot\vec{\mathsf{V}}(\mathsf{T}+\tau,\vec{X}_e(\mathsf{T})e^{\frac{\tau}{2}})\rangle}\delta(\mathsf{T}_1-\mathsf{T}_2) \qquad [61]$$

We solve [60] using [61] as an expression for $\frac{\delta\mathcal{A}}{\delta\vec{X}(\mathsf{T}_1)\delta\vec{X}(\mathsf{T}_2)}\left[\vec{h}=\vec{0}\right]$ and therefore, we deduce the second cumulant of $\mathcal{P}\left[\vec{X}(\mathsf{T})\right]$,

$$\langle\Delta\vec{X}(\mathsf{T}_1)\Delta\vec{X}(\mathsf{T}_2)\rangle = \overset{\leftrightarrow}{1}\, X_0^2 e^{\frac{\mathsf{T}_2+\mathsf{T}_1}{2}}\mathcal{J}\left(\vec{X}_0 e^{\frac{1}{2}\mathrm{Max}(\{\mathsf{T}_1,\mathsf{T}_2\})}\right)\left(1-\frac{\mathcal{J}\left(\vec{X}_0 e^{\frac{1}{2}\mathrm{Min}(\{\mathsf{T}_1,\mathsf{T}_2\})}\right)}{\mathcal{J}\left(X_0\right)}\right), \qquad [62]$$

with

$$\mathcal{J}\left(\vec{u}\right) = \frac{1}{|\vec{u}|^2}\int_0^\infty dt'\int_0^\infty d\tau\langle\vec{\mathsf{V}}(0,\vec{u}\,e^{\frac{\mathsf{T}'}{2}})\cdot\vec{\mathsf{V}}(\tau,\vec{u}\,e^{\frac{\mathsf{T}'+\tau}{2}})\rangle e^{-\mathsf{T}'-\frac{\tau}{2}}.$$

Using [45], $\mathcal{J}$ can be explicitly expressed in terms of position:

$$\mathcal{J}\left(\vec{u}\right) = \left\{\mathsf{K}_{\varphi_1\varphi_2}C_{\varphi_1}^{\varepsilon_1}C_{\varphi_2}^{\varepsilon_2}J\left(|\vec{u}|,\omega_{\varepsilon_1},\omega_{\varepsilon_2}\right)+\frac{4}{(1-\nu)^2}\mathsf{K}_{ss}c_s^2 J\left(|\vec{u}|,\omega_s,\omega_s\right)\right\} \qquad [63]$$

$$J(u,\alpha,\beta) = \frac{2}{\beta(\alpha+\beta-1)}\left\{\frac{\log(u)+\gamma_e-\frac{1}{2}-\frac{1}{2(\alpha+\beta-1)}}{u^2}+\frac{1}{u^{2(\alpha+\beta)}}\frac{\gamma_e(2\alpha+2\beta)}{2(\alpha+\beta)(\alpha+\beta-1)}-\mathcal{R}(u,\alpha,\beta)\right\},$$

where $\gamma_e$ is the Euler-Mascheroni constant, $\gamma_e(x)$ the Gamma function, and

$$\mathcal{R}(x,\alpha,\beta) = \int_x^{+\infty} du\left\{\frac{\left(\log(u)-\frac{1}{2(\alpha+\beta)-2}\right)(1+u)e^{-u}-\int_u^\infty dv v\log(v)e^{-v}}{u^3}+\frac{u^{-2(\alpha+\beta)-1}}{2(\alpha+\beta-1)}\int_u^{+\infty}dv\, v^{2(\alpha+\beta)-1}e^{-v}\right\}.$$

This function decays fast to 0 as $x>1$. Depending on the value of $\mathrm{Re}(\alpha+\beta)$, $J(u,\alpha,\beta)$ has for asymptotic trend $J(u,\alpha,\beta)\propto\frac{\log(x)}{x^2}$ or $J(u,\alpha,\beta)\propto\frac{1}{x^{2(a+b)}}$ which yields the asymptotic trends of the coefficient of variation $\mathrm{CV}(\mathsf{T})=\langle(\Delta\vec{X}(\mathsf{T}))^2\rangle^{1/2}/\langle\|\vec{X}(\mathsf{T})\|\rangle$ discussed in the main text. And for $\tau_c>2$, $\mathrm{CV}(\mathsf{T})\sim 2\mathrm{CV}_{\max}e^{-2\mathsf{T}/\tau_c}\sqrt{1-e^{-4\mathsf{T}/\tau_c}}$ as $X_0\gg1$ if the anisotropic feedback is low and oscillate around this trend if it is high.
